# A long-read RNA sequencing and polysome profiling framework reveals transposable element–driven transcript diversity and translational rewiring in glioblastoma

**DOI:** 10.64898/2026.04.18.719388

**Authors:** Mattia D. Pizzagalli, Shiven Sasipalli, Owen Leary, Lily Tran, Brian Haas, Nikos Tapinos

**Affiliations:** Laboratory of Cancer Epigenetics and Plasticity, Brown University, Providence, RI, USA; Methods Development Laboratory, Broad Institute of MIT and Harvard, Cambridge, MA, USA; Department of Neurosurgery, Brown University Health, Providence, RI, USA; Tinos Therapeutics Inc., Cambridge, MA, USA

## Abstract

**Background:** Transposable elements (TEs) account for over half of the human genome and are often derepressed in cancer. TEs can add cryptic splice sites, undergo exonization, and generate gene–TE fusion transcripts, but the combined effects of TEs on RNA processing and translation in glioblastoma stem cells (GSCs) remains incompletely elucidated.

**Results:** We combined long-read RNA sequencing with polysome profiling in four patient-derived GSCs and two neural stem cell (NSC) controls to resolve TE-associated transcript diversity and its relationship to ribosomal engagement. Across GSCs, we identified 13,421 alternative splicing (AS) events, 3,077 of which contained TEs within 150 bp of splice junctions. AS sites proximal to TEs were associated with increased isoform switching compared to non–TE-associated AS sites (odds ratio 2.9 - 4.3). Moreover, AS isoforms generated from TE-proximal sites were more likely to exhibit altered ribosomal association (odds ratio 2.54). Directional shifts were observed, with shorter isoforms associating with monosome fractions and longer isoforms with polysome fractions. To enable systematic detection of gene - TE chimeric transcripts, we developed FuTER (Fusion TE Reporter), a long-read–based framework for identifying TE-associated fusions. Application to GSC datasets identified 78 GSC enriched fusion transcripts, several supported by breakpoint-spanning reads in polysome fractions, consistent with ribosome association.

**Conclusions:** Our data suggest that TEs correlate with abnormal splicing activity and altered ribosome engagement in glioblastoma stem cells. By integrating long-read sequencing with polysome profiling and fusion detection, we establish a framework for analysis of TE-induced transcript diversity and its effects on cancer evolution and plasticity.

## Introduction

Transposable elements (TEs) are a diverse group of mobile genomic entities that can reinsert themselves into new genomic loci (1). TEs account for more than half of the human genome and contribute significantly to the genomic diversity and architecture of cells, as well as the regulation of gene expression (1–4). Under normal conditions, human cells employ robust epigenetic and post-transcriptional mechanisms to suppress TE expression and prevent their potential deleterious effects (5). However, these mechanisms can be weakened in disease states, particularly cancer, leading to TE reactivation and widespread transcriptional consequences (6–8).

There are two main classes of TEs defined by their method of reinsertion. Class I retrotransposons propagate using an RNA intermediate in a copy-and-paste mechanism involving reverse transcription and then reinsertion into the genome (1). Class II DNA transposons replicate via a cut-and-paste mechanism, which is driven by transposase activity to reinsert sequences into a new location (1). Retrotransposons, which dominate the human genome, include Long Terminal Repeat (LTR) containing elements and non-LTR containing retrotransposons. Non-LTRs contains families such as the Long Interspersed Nuclear Elements (LINEs) and Short Interspersed Nuclear Elements (SINEs), each of which represent 17% of the human genome (1). LTR retrotransposons consist primarily of endogenous retroviruses (ERVs), which account a further 8% of the human genome (1). Although these elements are ancient remnants of past insertions and are often no longer mobile, many retain regulatory potential or can influence transcriptional programs through insertional mutagenesis, promoter activation, and RNA processing effects (1,9–12).

Aberrant reactivation of TEs has emerged as a hallmark of several cancers, where hypomethylation and chromatin remodeling result in elevated TE transcription and mobilization. This can promote oncogenesis through multiple mechanisms, including increased genomic instability, interference with gene regulation, activation of immune signaling, driving expression of alternative isoforms or producing novel fusion transcripts (6–8,10). Despite the growing recognition of TE dysregulation in malignancies, the mechanistic contributions of TEs to transcriptomic diversity remain incompletely understood, particularly in highly heterogeneous and aggressive tumors like glioblastoma (GBM).

GBM is the most common malignant primary brain tumor, characterized by rapid progression, therapy resistance and a five-year survival rate of 6.9% (13,14). Within these tumors, glioblastoma stem cells (GSCs) represent a subpopulation with self-renewal capacity and tumor-initiating potential, thought to underlie therapeutic resistance and recurrence. GSCs exhibit extensive epigenetic reprogramming, transcriptional plasticity, and genomic instability, features that may be influenced or amplified by TE activity (15,16). Indeed, previous studies have demonstrated elevated expression of ERV transcripts and the presence of hundreds of TE-derived peptides unique to GBM (17,18). However, the extent to which TEs contribute to fusion transcript formation or alternative splicing in GSCs remains unknown.

Chromosomal instability (CIN) is a defining feature of GBM and other cancers, promoting structural rearrangements that drive oncogenic evolution (19–21). In cancer, increases in CIN contribute to genomic aberrations, deepening the genomic heterogeneity of tumors and promoting the survival and proliferation of cells that are conferred a survival advantage from changes in genes or regulatory pathways (19,21). One major consequence of CIN is the formation of fusion transcripts, which are hybrid RNA molecules derived from two distinct parental genes (22). These chimeric transcripts can lead to novel protein products, altered expression of parental genes, or generation of neoantigens that influence tumor immunogenicity (22,23). It has been suggested that fusion transcripts contribute to the development of 16.5% of cancers cases (and are the sole driver mutation in more than 1%) (23). Although many fusion transcripts arise from genomic rearrangements, transcriptional read-through and splicing between adjacent loci can also produce fusion RNAs independently of DNA-level changes (22). TEs have been implicated in both mechanisms, as they can mediate recombination events that lead to rearrangements or supply cryptic splice donor and acceptor sites that facilitate RNA-level fusions (24–28).

Beyond their role in fusion transcript formation, TEs also serve as a vast reservoir of alternative splice sites and regulatory sequences. Alternative splicing enables a single gene to produce multiple mRNA isoforms with distinct functions (29). In cancer cells, dysregulation of the splicing machinery, along with global genomic hypomethylation and altered transcription dynamics, can activate cryptic splice sites within intronic TEs, resulting in the inclusion of TE-derived exons or exonized fragments (30,31). This phenomenon, particularly enriched in Alu elements, expands the proteomic and functional diversity of cancer cells. In GBM, aberrant splicing of key regulators such as ITSN1 has been linked to enhanced aggressiveness and invasion, suggesting that alternative splicing may play an unrecognized role in defining malignant phenotypes (32,33).

Although multiple computational frameworks exist to detect TE-related fusion transcripts using short-read RNA-seq data, these approaches are limited by their inability to resolve full-length transcripts or accurately identify fusion junctions (34,35). The advent of long-read sequencing technologies provides an opportunity to overcome these challenges, as full-length reads can span complete transcripts and delineate exact fusion breakpoints between TEs and their partner genes. A recent study leveraged this advantage to more accurately identify fusion transcripts arising from genes and TEs (36). However, this approach utilizes the existing Gencode and Repeat Masker annotations, limiting their ability to identify fusions arising from novel or un-annotated TE insertions. In the present study, we developed FuTER (Fusion TE Reporter) as a computational framework to sensitively and specifically identify TE–gene fusion transcripts from long-read data (37). Using simulated datasets, we benchmarked its precision and recall, demonstrating high performance across a range of expression levels and insertion types. We then applied FuTER to a library of long-read transcriptomes we generated from patient-derived GSCs to systematically profile TE-driven fusions and exonization events. This allows us to comprehensively identify TE-gene fusion transcripts, including those arising from novel insertions.

Our study presents the most comprehensive analysis to date of how TEs reshape the transcriptome in glioblastoma stem cells. By integrating long-read sequencing with a refined computational workflow, we reveal that TEs contribute to both structural and regulatory transcript diversity in GBM. These findings establish a methodological framework for TE fusion discovery and offer new insights into the molecular mechanisms driving transcriptomic complexity and tumor recurrence in glioblastoma.

## Results

### Alternative Splicing in GSCs vs NSCs

To most effectively study the dynamics of alternative splicing in GSCs, we performed long read sequencing of four different patient derived GSCs and two neural stem cell (NSC) lines to a depth of at least 30 million reads (avg: 42 million reads, **Supplementary Table 1**). This increased depth allowed us to identify rare fusion events that are sometimes missed by long read sequencing approaches and to our knowledge represents the only long read RNA sequencing library of GSCs currently available. We utilized Bambu followed by SQANTI to generate a high confidence transcriptome across the cell lines, which we used to perform a genome-wide analysis of splicing patterns and isoform usage (**Figure 1a**) (38,39).

**Figure 1:**
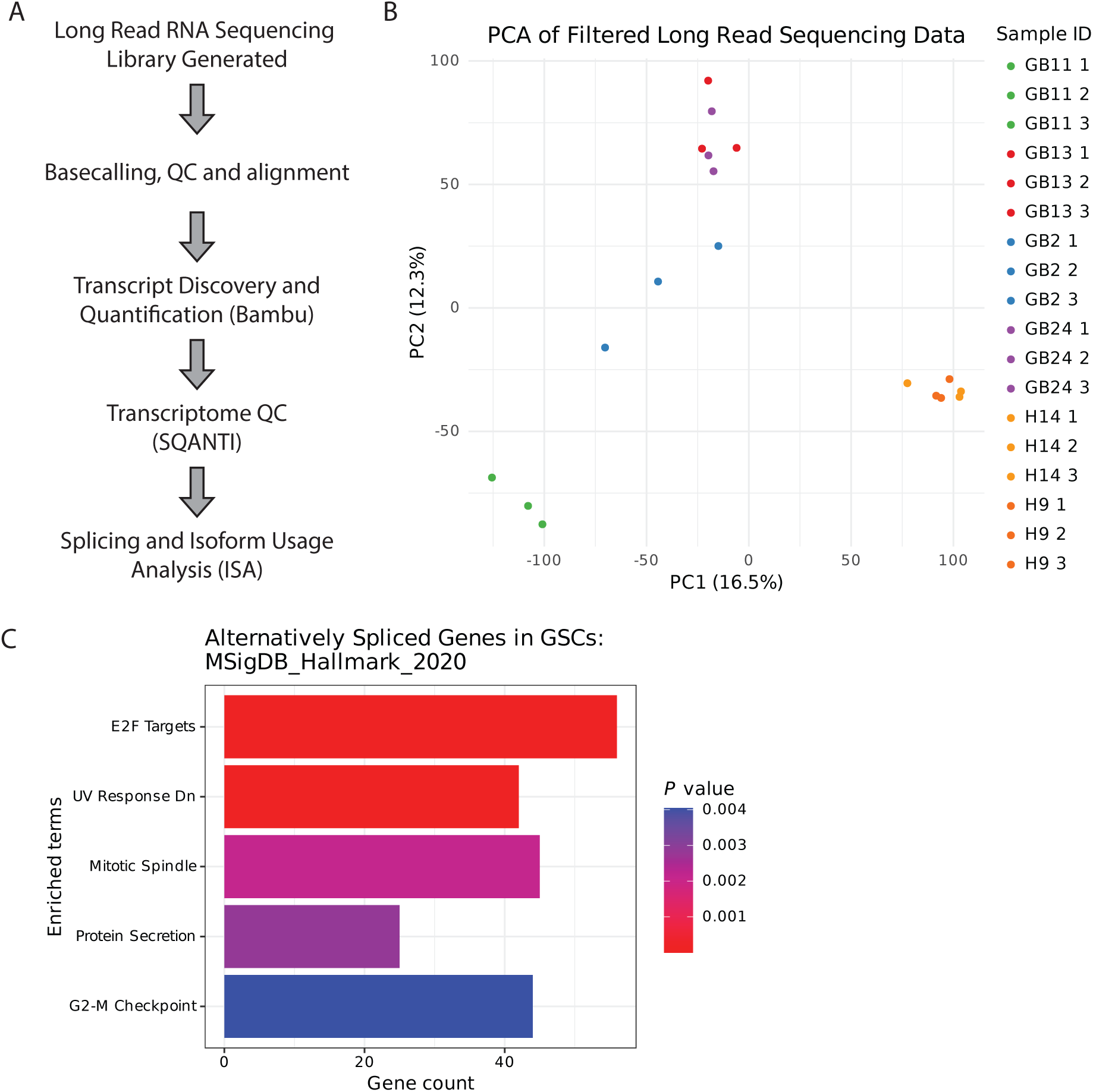
Using Long Read Sequencing to Study Alternative Splicing in GSCs. **(A)** A schematic of the approach used to study alternative splicing and isoform usage changes in GSCs. **(B)** The PCA plot of transcript expression in GSCs. The NSCs cluster very tightly (orange points) while GSCs demonstrate the heterogeneity that defines these plastic cells. **(C)** Genes that are alternatively spliced in GSCs enrich for critical cancer hallmark pathways like the G2-M Checkpoint, Targets of E2F and down regulators of UV damage response.

As expected, initial analysis of these datasets demonstrated that the NSCs clustered tightly together while GSCs demonstrated more variance in their transcriptomic profiles (**Figure 1b**). Due to the heterogeneity of the GSC samples, differential splicing analysis was performed for each GSC independently against the NSC samples. Our analysis demonstrated that GB2s have a splicing profile most like NSCs, with 3969 total alternative splicing events identified. Analysis of GB11, GB13 and GB24 identified 6,765, 6,756, and 6,823 AS events respectively. Across all four GSCs, the most common changes to transcript structure were alternative transcription start sites and termination sites (**Supplementary Figure 1**).

To understand the effect that these splicing patterns have on the transcriptome of the cells, we also studied isoform usage changes in the cells. This allows us to identify when there is a significant change in the expression distribution of a given gene that results from AS. Underscoring the importance of splicing patterns to the identify of cells, the genes impacted by AS in all four GSCs enriched for critical cancer hallmark pathways like E2F targets and the G2M checkpoint, and included genes central to GSC identify like FOXM1, ECT2, CDC25A and CDCA7L (**Figure 1c**).

### Involvement of TEs in Alternative Splicing

To identify the role TEs play in the dynamics of alternative splicing, we focused on AS events that overlapped or were within 150 bp of a TE. Recent studies have shown that TEs inserted within 150 bps of an exon border have a significant regulatory impact on splicing (40,41). Between the NSCs and the GSC samples we identified 13,421 unique AS events, 3077 of which were identified to have proximal TEs. These overlaps arose from 2148 different TEs (519 LINE, 1078 SINE, 280 LTR, 271 DNA). Across all four GSCs these splicing events had a similar distribution of TE association suggesting that these effects are not cell line specific but rather that TEs have a consistent role in splicing across cell lines (**Figure 2a, Supplementary Figure 2**). To study whether a TE being proximal to a splicing site is correlated to altered isoform usage, we identified transcripts that had isoforms with or without TEs proximal to their splice sites and investigated rates of isoform switching. In all four GSCs, genes undergoing AS with proximal TEs were significantly more likely to have isoform switching. Whereas 21% of genes without a TE proximal AS event have isoform switching in GB24, genes with a TE isoform switched 51% of the time (Odds Ratio: 3.94 [3.43-4.55] (p-value: 2.2e-16). GB2 had an odds ratio of 2.88 [2.45-3.38] (p-value: 2.2e-16) while GB11 and GB13 had odds ratios of 3.82 [3.32-4.41] (p-value: 2.2e-16) and 4.28 [3.71-4.92] (p-value: 2.2e-16) respectively (**Figure 2b**). These results support our hypothesis that TEs play a significant role in regulating splicing and isoform usage in GSCs and thereby contribute to the identity of these cells. We next identified isoforms whose usage differed significantly between a GSC and NSC and had unique TE overlaps at the splice sites of the isoforms (eg. a TE overlaps with the splice site that differentiates the isoforms). These represent cases where the TEs may directly contribute to the switch through their regulatory roles in splicing. Across the four GSCs, there were 1,135 AS events resulting in an isoform switch with a TE proximal. Representative examples relevant to glioblastoma biology and pathology are shown in **Figure 2c**. For H2BC15, the isoform switch results in an increased expression of a protein coding isoform, while the isoform expressed evenly between GSCs and NSCs (ENST00000449538.3) results in likely nonsense mediated decay. H2BC15 is a recently discovered novel histone variant that may bestow unique properties to nucleosomes, suggesting these TE driven isoform switches may be altering the regulatory landscape of the GSCs (42). PAK4 isoform switching results in an increased expression of a different protein coding isoform (ENST00000599386.5), in addition to the canonical isoform (ENSG00000360442.8) that is expressed evenly between the GSCs and NSCs.

**Figure 2:**
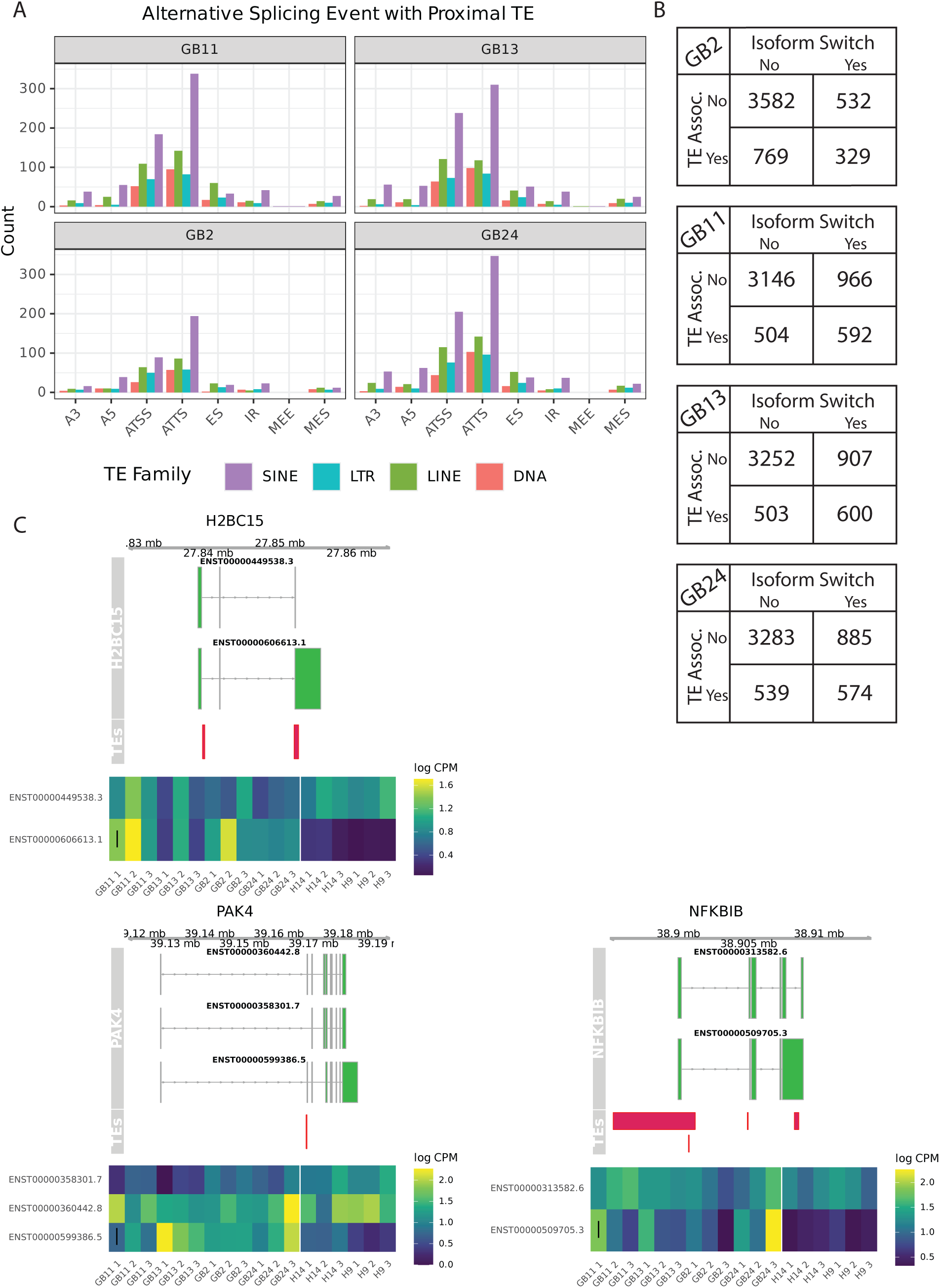
Transposable Element Linked Alternative Splicing Alters Isoform Usage in GSCs. **(A)** 3077 alternative splicing events occur with a TE within 150 bp of the splice site. These events are shown separated by the family of the associated TE. SINE elements are the most commonly associated family and often are found proximally to alternative transcription termination sites. We also found that LINE elements are primarily associated with alternative transcription termination and alternative transcription start site events, findings which support previous studies. **(B)** Transcripts containing a transposable element associated splice site are significantly more likely to undergo isoform switching, suggesting transposable elements play a critical role in shifting the profile of expressed isoforms. **(C)** Representative demonstrations of genes with significant isoform switches involving a transcript(s) with transposable element proximal alternative splicing events. The top panel displays the structure of the transcripts, as well as the overlapping transposable element(s). The bottom panel shows expression levels across all samples. The horizontal black line indicates which isoform has increased usage in the GSCs. H2BC15 switches from an isoform that undergoes nonsense mediated decay to a protein coding isoform containing an alternative transcription termination site. The splice site overlaps with an AluSX element. PAK4 expression shifts towards isoforms containing an exon whose alternative 5’ splice site overlaps with a L2 element. NFKBIB has an increase of an isoform with an intron retention event that overlaps with an AluSg element.

### Identification of Fusion Transcripts Involving Transposable Elements

After studying the involvement of TEs in alternative splicing dynamics, we sought to specifically identify chimeric gene-TE transcripts. These can arise from both the exonization of TEs into an existing transcript due to the cryptic splice sites present in TEs or from either genome dependent or independent fusion processes. To this end, we developed FuTER (Fusion TE Reporter), a pipeline dedicated to identifying gene-TE chimeric transcripts from long read sequencing data and the first that can identify fusions arising from novel TE insertions (**Figure 3a**). Our approach was based on CTAT-LR-Fusion, an existing gene fusion identification pipeline, which we adapted here for detection of transcripts involving fusions of TEs with known human genes (37). The process of identifying fusion transcripts involving TEs is composed of four main phases. First, a library of reference TEs was generated using RepeatMasker. To focus on TEs most likely to contribute to fusion transcripts, we filtered for Class I and II TEs with a calculated divergence less than 25%. This removes ancient elements that, due to an accumulation of mutations, are unlikely to either maintain their ability to reinsert in the genome or contain functional open reading frames. This cutoff can easily be changed by users for their use cases. In our case, we generated a TE reference library containing 1.6 million elements (See **Supplemental Table 2** for an overview). Long reads are aligned to this library and those that align to a TE are passed to the next phase, where reads are aligned to a combined genome containing the human genome and the library of TE sequences. To prevent alignment to genomic TE sequences, all non-exonic regions of the genome annotated by RepeatMasker as TE-derived are masked. This ensures that segments of reads derived from TEs are readily identified through their alignment to a TE library sequence. This is particularly important for TE exonization events, which otherwise would not be as easily identified. This alignment step utilizes a customized version of minimap2 developed for CTAT-LR-Fusion that only reports reads that align to multiple loci. Next, gene and TE partners of candidate fusion transcripts are identified based on the loci that chimeric reads align to in the combined genome/TE database. In the third phase, the full-length sequences of the gene and the TE of candidate loci are extracted and combined to create a new fusion contig. Reads are then realigned to these contigs and the specific breakpoints are identified based on a minimum number of high-quality alignments. For more details, see Methods and Haas et al (43).

**Figure 3:**
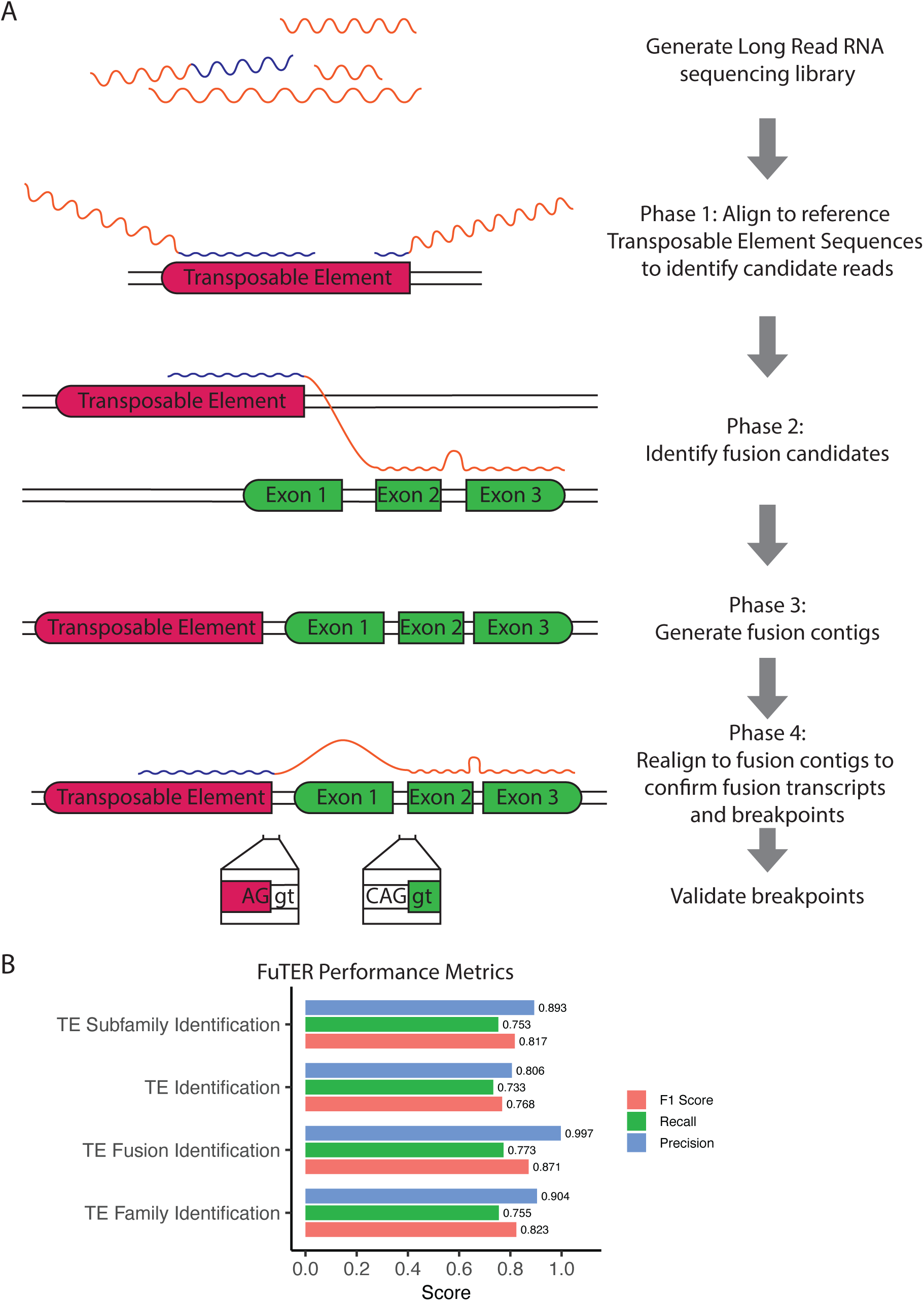
Schematic and Benchmarking of Fusion TE Reporter (FuTER) **(A)** A schematic depicting the approach used by FuTER. **(B)** Benchmarking of FuTER using a simulated long read library containing normal transcripts, gene-gene fusions and gene-TE fusion transcripts demonstrates strong performance. Performance was tuned to maximize precision and minimize the risks of false positives.

### Evaluation of Performance Using Simulated Reads

To evaluate the pipeline’s performance, we simulated three libraries of long reads that contained 2,500 reads derived from 500 gene-TE fusion transcripts, 12,500 reads derived from 2,500 gene-gene fusion transcripts and all annotated transcripts in the human genome (**Supplementary Table 3**). This wide-ranging approach allowed us to evaluate the pipeline’s precision and sensitivity in identifying TE-gene chimeras across different levels of granularity. The pipeline was evaluated for its ability to identify gene-TE chimeras across the three simulated libraries, as well as how accurately it identifies the individual components of the simulated fusions. Our approach demonstrated strong performance (F1 score: 0.871) with particularly high precision (0.997) (**Figure 3b**). Although recall is lower (0.773), tuning performance for higher precision allows researchers to begin validation studies with more confidence. The pipeline retained strong performance in terms of identifying the exact partners involved in a TE-fusion transcript (Precision: 0.806, Recall: 0.733, F1 Score: 0.768; **Figure 3b**). One notable aspect is that there is a degree of variance in the pipeline’s performance between the different TE families as well as the length of the TE component in the simulated fusion (**Supplementary Figure 3**). Overall, our results suggest that very short TE components are often not identified, presumably due to the challenges in identifying unique patterns that are used by the minimap2 aligner to map a read to a specific region of the genome. On the other hand, across all families of TEs, TE partners that were misclassified according to TE family tend to be longer. Overall, our benchmarking shows that the pipeline identifies TE-gene chimeric transcripts with a very high degree of precision and strong recall.

### Analysis of HG002 and HCC1395 direct RNA sequencing data

To further validate our approach, we applied FuTER to a set of long read sequencing results from cell lines with publicly available genome assemblies. By analyzing data from cell lines with a fully annotated genome, we sought to identify TE fusion reads from cell lines with genomic evidence of novel TE insertion events to demonstrate the utility of our approach. We were able to identify gene-TE chimeric transcripts in both cell lines, HCC1395 and HG002 (**Figure 4**). In the HCC1395 samples, we identified a fusion involving KMT2E and AluSx that demonstrates alignment with a TE element not found in the standard genome but identified ∼105kbp away in the HC1395 genome. Interestingly, this transcript skips over multiple AluSx insertions more proximal to KMT2E, suggesting a more targeted event may give rise to this fusion transcript (**Figure 4a**). Furthermore, manual inspection of the breakpoints within the gene and TE clarify that the reads align to consensus splice motifs. We also identified a fusion transcript derived from PTEN and a LTR47A insertion located 44 kbp away in the HC1395 genome that also contains consensus splice sites at the predicted breakpoints between the gene and the TE (**Figure 4b**). A similar TE is also identified in the standard human genome, suggesting it is not a novel insertion. In HG002, two fusion transcripts were identified by the pipeline. The first is a fusion transcript containing ZNF732 and an L1MA6 moiety identified by a set of four reads (**Figure 4c**). The other, involving ESRG, had a high rate of insertions and deletions from the reads that mapped to the fusion, indicating potential poor read quality or mapping (**Supplementary Figure 4**).

**Figure 4:**
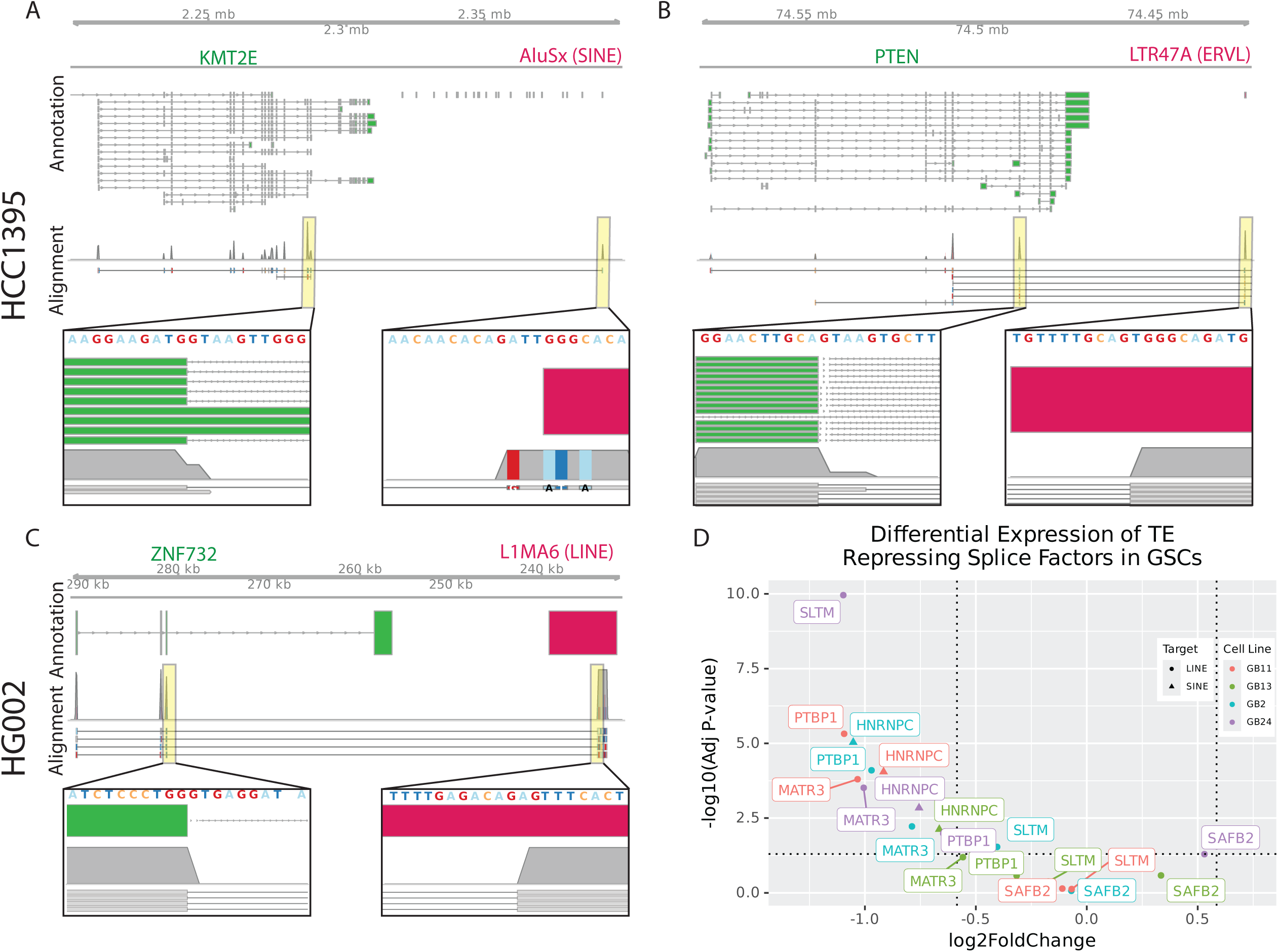
Validation of FuTER in HCC1395 and HG002 Sequencing Data and Altered Expression of Splice Regulators in GSCs. **A-C)** Figures depicting gene-TE fusion transcripts identified by FuTER in HCC1395 (A, B) and HG002 (C) direct RNA sequencing data. Top panel (Annotation) of figures shows the structure of the gene (in green) and the TE (in red). The bottom panel (Alignment) shows alignment of reads to the region of the genome giving rise to the fusion transcripts. Inserts show the sequences and alignments in more detail surrounding the breakpoint. Two of the three fusions have more than four reads that cross the fusion breakpoint. All three identified fusions align consistently with gene structures. **(D)** Differential expression analysis of GSC expression data demonstrated significant changes in the expression of splice regulators that typically inhibit the exonization of TEs. HNRNPC, which is significantly downregulated in all four GSCs, competitively inhibits the U2AF65 splicing factor by binding to Alu elements. MATR3 and PTBP1 form a complex that inhibits LINE exonization. Both proteins are significantly downregulated in GB2, GB11 and GB24.

### TE Gene Fusions in Glioblastoma Cells

After establishing FuTER’s ability to identify gene-TE fusion transcripts, we sought to characterize the landscape of these structures in our GSC and NSC long read RNA sequencing samples. Because TE exonization events are typically repressed in cells, we first investigated whether regulatory proteins that typically prevent TE exonization events are differentially expressed in GSCs (**Figure 4d**). Across all four GSCs, HNRNPC, a splice regulator that prevents exonization of SINE elements, is significantly down regulated (Fold Change > 1.5, Adj p-value < 0.05) (44). GB2, GB11 and GB24 all also demonstrate significant down regulation of MATR3 and PTBP1, which together inhibit the exonization of LINEs (45). The changes in the expression of these regulatory elements, amongst others, suggests these cells are primed for the exonization of TEs. FuTER identified a series of 78 TE-fusions enriched in GSCs (**Table 1**). Overall, fusions arose from genes found throughout the genome, without any enrichment for specific regions. These transcripts involved all four families of TEs we explored, including 11 DNA elements, 20 LINEs, 13 SINEs, and 35 LTR elements. Interestingly, we identified 12 fusion transcripts where a TE initiated transcription and 66 fusions beginning with a gene transcript and terminating in a TE. Because our benchmarking did not demonstrate a difference in performance between the identification of Gene-TE versus TE-Gene fusions, we consider that this observed variability in TE-positioning within TE-fusion transcripts is due to the underlying biology. One possibility is that gene transcription start sites are more efficient at initiating the production of transcripts to a level where we can identify them.

**Table 1:**
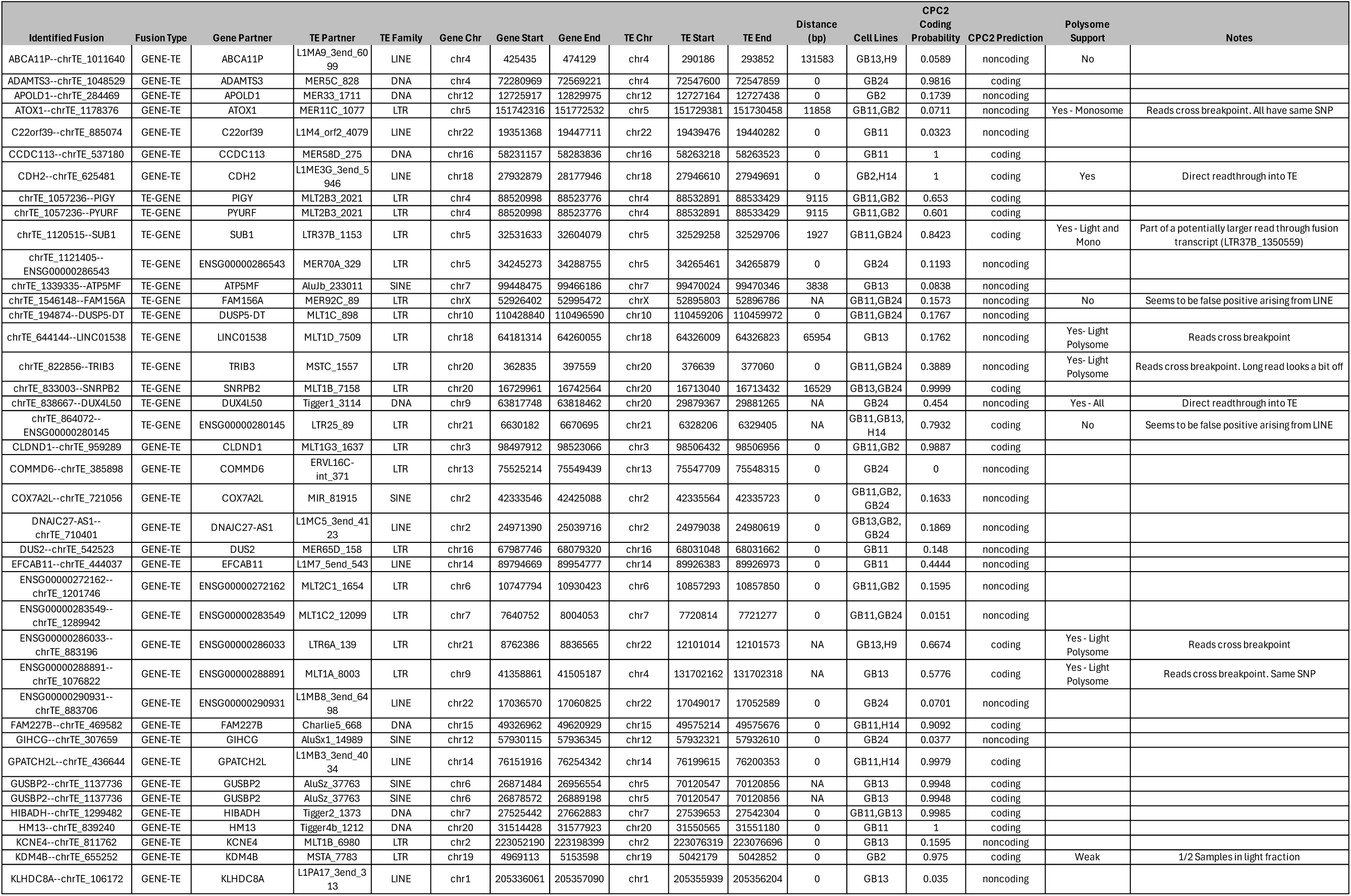
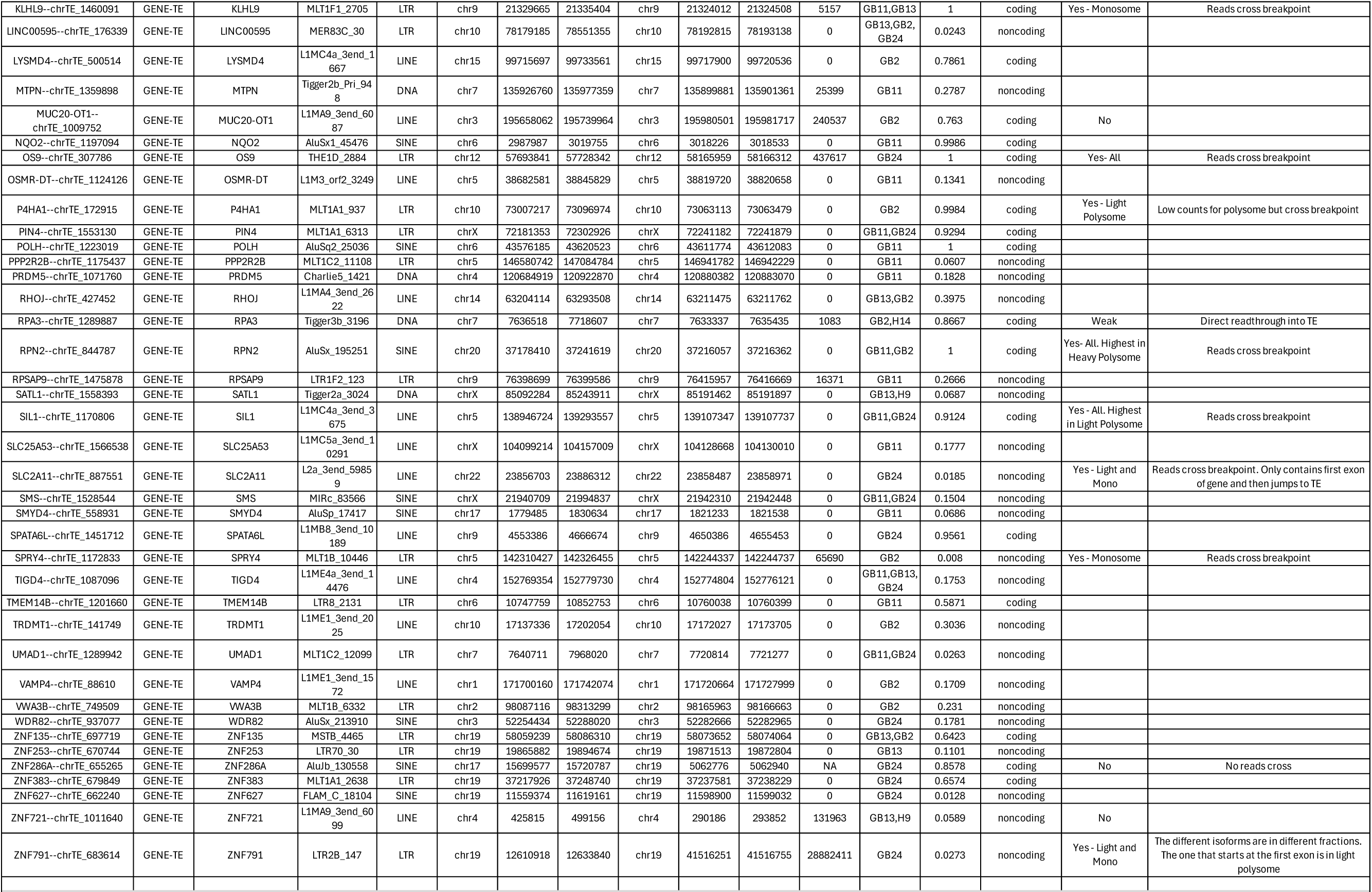
A summary of all TE gene fusions identified and enriched in GSCs. The numerical value at the end of the TE Partner values are unique identifiers that allow us to map TEs back to their point of origin

Evaluating the distance between genes and the identified TE partner, FuTER identified 54 fusions between a gene transcript and an internal TE, underscoring this approaches ability to identify TE exonization events. Of the 24 remaining fusions, 5 identified potential chromosomal rearrangement events (or fusions arising from novel TE insertions) and the rest represented fusion transcripts between a gene transcript and a downstream TE ranging from 1083 bp to 28,882,411 bp away (mean: 1,768,009 bp, median: 16,529 bp). We identified fusions involving known oncogenes such as RPN2, SPRY2 and P4HA1, as well as other cancer-related genes like ABCA11P, ZNF791 and ATOX1. We also identified 21 fusions between non-coding RNAs and TEs. Similar structures have been reported previously in literature and have functional roles in cells (28).

We next studied the individual breakpoints between the genes and TEs in more detail to ensure the breakpoints represented potential splice sites and aligned well to gene structures. Representative figures are shown in **Figure 5** and **Supplementary Figure 5**. Overall, most fusions that were identified conformed to gene exon structures and their TE breakpoints contained either consensus splice motifs or motifs that were identified by SpliceAI as potential cryptic splice sites (46). We also investigated the coding potential for the fusion transcripts using CPC2 to study whether the open reading frames are preserved (47). Of the 78 different fusion transcripts identified, 34 (43.6%) were identified to have a complete ORF and classified as a coding sequence (**Table 1**).

**Figure 5.**
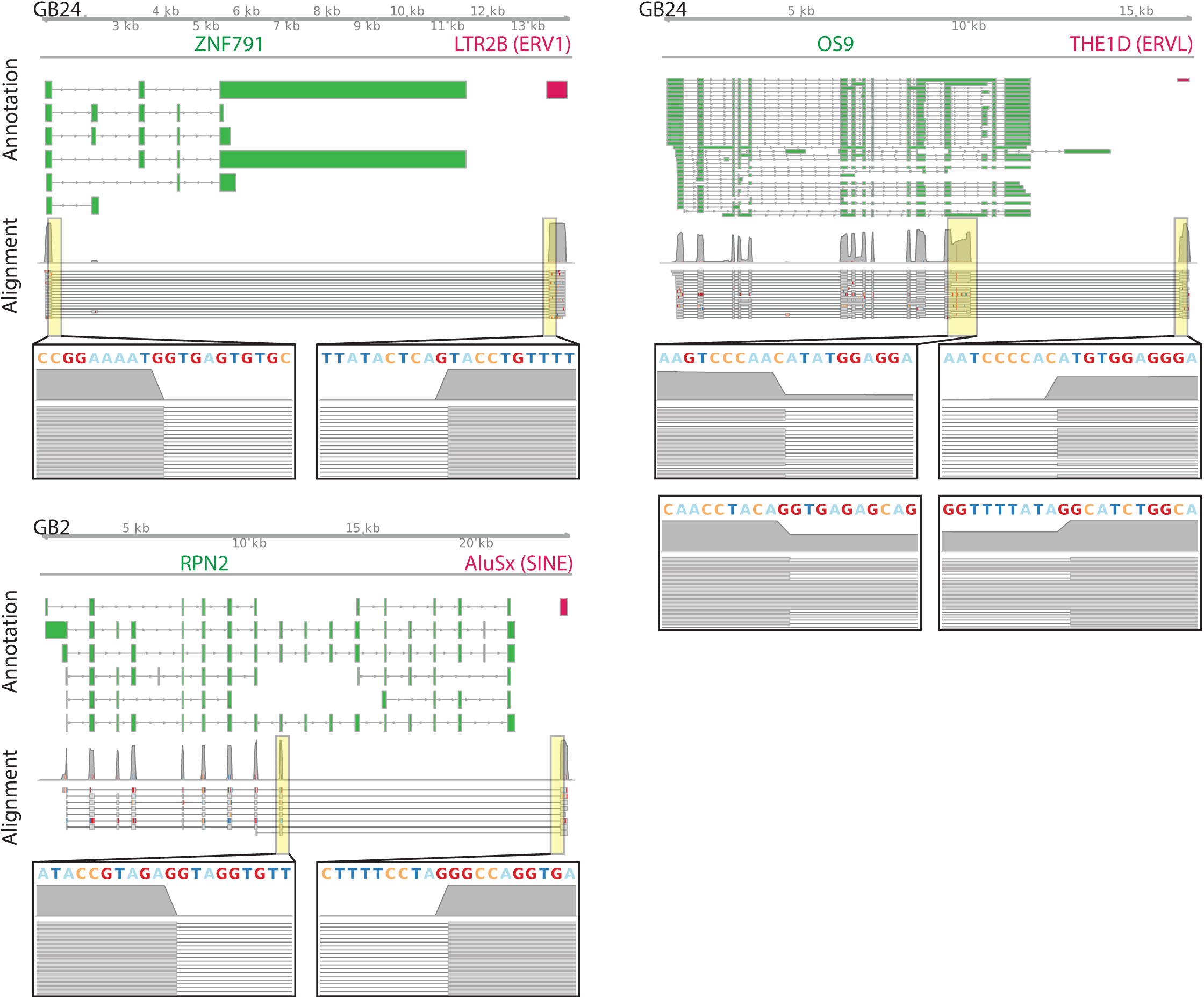
FuTER Identified Fusion Transcripts. Representative images of reads spanning fusion transcript breakpoints of fusion transcripts identified by FuTER in the indicated GSC. Reads were extracted and aligned to fusion contigs generated by FuTER. Identified fusions typically conform to gene structures up to the breakpoint and contain consensus splice sites or predicted cryptic splice sites on the TE side of the fusion transcript. Fusion structures are depicted in the top panel (Annotation) while long reads are shown below (Alignment). Inserts depicted the exact sequences surrounding the breakpoint to visualize splice sites and breakpoint precision. Multiple reads in independently generated samples cross the fusion break point and support its existence. The RPN2-AluSx fusion transcript was also identified in GB11 with the same alignment pattern.

### Polysome Profiling Demonstrates Association of Fusion Transcripts with ribosomes

After interrogating the involvement of TEs in alternative splicing and chimeric transcripts, we sought to identify whether these patterns are replicated in the translating ribosomal fractions of RNA. To this end, we performed polysome sequencing of three of the GSC lines, GB2, GB24 and GB13. By quantifying changes in each isoform’s ribosome occupancy (monosome, light polysomes and heavy polysomes) as a fraction of the gene’s total ribosome bound expression, we can infer which transcript structures are associated with efficient ribosome engagement. Although polysome profiling does not provide definitive proof of translation, association of a transcript with heavy polysomes has repeatedly been shown to result in increased translation while monosome associated transcripts are generally less efficiently translated (48–51). A notable exception to this is shorter transcripts, which can still be translated from monosomes (52). Although ribosomal stalling or shifts of ribosomes towards untranslated open reading frames can occur and cloud the interpretation of polysome profiling, the initiation of translation (ie the association of transcripts with ribosomes) is thought to be a significant source of translational regulation and association with heavy polysomes is understood to typically result in translation (49–51,53). Our analysis demonstrated that within the heavy polysome fraction, isoforms of genes without an alternative transcription start site (ATSS) have a much higher usage than in the light and monosome fractions, implying that these gene isoforms could be translated more efficiently. This pattern is also replicated for isoforms with an alternative transcription termination site (ATTS) (isoforms with an ATTS represent a smaller fraction of a gene’s overall expression in the heavy polysome versus light polysomes or monosomes) (**Figure 6a**). Across the three GSCs, we also identified that isoforms with an intron retention event comprise a lower overall proportion of a gene’s expression while isoforms without the intron retention have their relative usage increase, implying that there is preferential ribosome loading of canonical isoforms.

**Figure 6.**
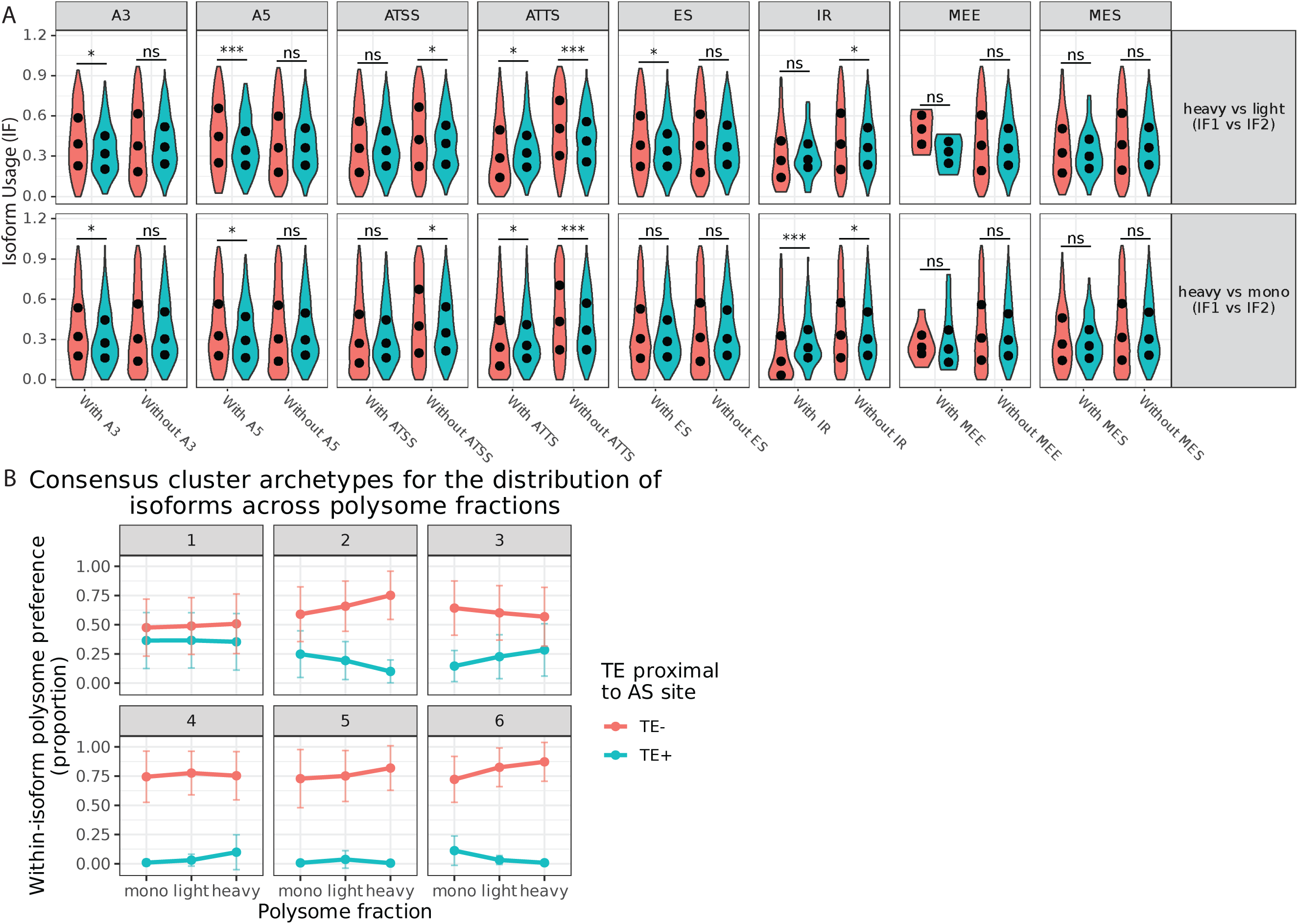
Analysis of Alternative Splicing in GSCs. **(A)** Alternative splicing analysis of GSCs followed by quantification of isoform switching across polysome fractions demonstrated that alternative splicing impacts the association of isoforms with polysomes. Isoforms that utilize canonical transcription start and termination sites are significantly enriched in the heavy polysome fraction versus both light polysome and monosome fractions when compared to isoforms utilizing alternative start and termination sites. Isoforms that have an alternative 5’ or 3’ splice site also contributed a significantly larger fraction of a gene’s expression on heavy polysome fractions versus other fractions. **(B)** Consensus clustering demonstrated that genes expressing isoforms both with (TE+) and without TEs (TE-) proximal to their splice sites form six stable clusters. These clusters represent distinct profiles of isoform fractions contributed by either TE+ or TE- isoforms to overall gene expression. Cluster 3, which consists of genes with an increasing proportion of their expression arising from TE+ isoforms, contains multiple genes associated with mTORC1 signaling, the p53 pathway and the G2-M checkpoint, suggesting these critical pathways may be affected by alternative splicing influenced by TEs.

To identify how TE insertions proximal to alternative splicing sites effect ribosome association of transcripts, we studied the proportion of a given isoform’s expression across the monosome, light and heavy polysomes. Using consensus clustering approaches, we identified six general patterns of TE isoform expression (**Figure 6b**). Cluster 1 genes have a more even distribution of expression across the ribosomal fractions arising from isoforms with and without TEs proximal to AS sites, while Clusters 2-6 express isoforms without TEs influencing AS events at higher proportions.

Clusters 2-6 differ from one another with regards to the pattern of ribosomal affinity depending on the presence of TEs proximal to the AS sites of the different isoforms. Genes in clusters three and four have an increased fraction of their overall expression arising from the TE associated splicing events, suggesting that the likely translated transcripts have an increased proportion of these alternative transcripts (**Figure 6b**). Cluster 3, which has the largest gain in the expression of these TE associated isoforms includes multiple genes associated with mTORC1 signaling, the p53 pathway and the G2-M pathway, suggesting that the translational efficiency of genes associated with these processes central to cancer cell proliferation may be altered by changes in splicing caused by TEs.

We also hypothesized that genes that have isoforms with TEs located proximal to AS sites are more likely to alter the expression of isoforms between the heavy polysome and monosome fractions. We found that TE presence proximal to a splice site strongly increases the odds of isoform switching, even when controlling for gene variability and transcript length (Odds ratio 2.54 [1.9 – 3.4], p value: 3.6e-10). Within genes, shorter isoforms preferentially switch to monosomes (Odds ratio per doubling of transcript length ∼0.71 [0.66-0.77], p value: 1.62e-16) while longer TE+ isoforms shift towards heavy polysomes (Odds ratio per doubling of transcript length ∼1.26 [1.17-1.37], p value: 9.35e-9). Although a shift towards the monosome for shorter isoforms can be explained by both inefficient or stalled translation due to early termination sites, shorter transcripts have also been shown to translate from monosomes, suggesting these switches may still result in alternative protein products. On the other hand, the shift towards the heavy polysomes for longer transcripts provides strong evidence that alternative splicing products related to TE insertions are efficiently translated. Altogether, these data show that the presence of a TE proximal to a AS site increases the dynamism of affected transcripts and contributes to changes in expressed and potentially translated isoforms.

### Association of TE-fusion chimeric transcripts with translating ribosomes

We next used the polysome sequencing data to both validate the existence of our identified gene-TE chimeric transcripts and identify whether they engage with ribosomes. For fusion transcripts arising from potential exonization or readthrough events, we aligned reads to the human genome and sought to identify reads that cross the same breakpoints identified by long read sequencing. Of the seven exonization/readthrough fusion transcripts we investigated, six contained reads that cross the same breakpoints identified using FuTER. For fusion transcripts arising from genes and TEs separated by greater distances, we aligned reads to a combined genome of the fusion contigs and the human genome and focused on primary alignments to the fusion contigs, an approach previously utilized by FusionInspector (54). Of the 24 gene-TE fusion transcripts investigated, 18 fusions had multiple reads cross the breakpoint in the polysome fractions. The OS9-THE1D(LTR) fusion transcript had multiple reads cross the breakpoint across all ribosome fractions, suggesting that this fusion transcript efficiently binds to ribosomes and potentially is translated (**Figure 7**).

**Figure 7.**
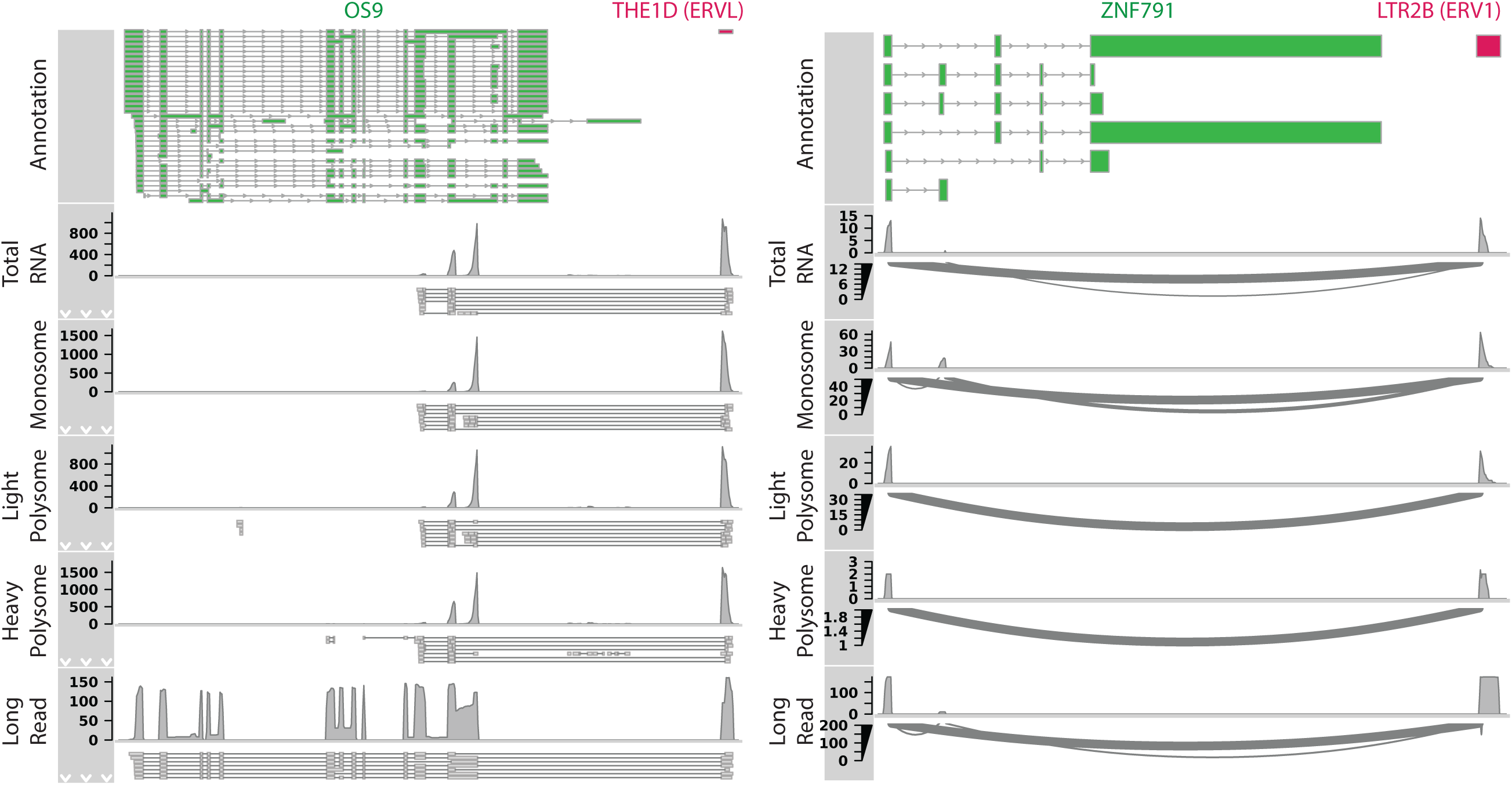
Polysome Profiling Supports Validity of FuTER Identified Fusion Transcripts. Polysome profiling further supported the existence of several identified fusion transcripts (11 of the 17 fusions we manually investigated). Figures depict the structure of the fusion contig in the top panel (Annotation), followed by the four short read RNA seq samples derived from the polysome pro ling samples. Reads that primarily align to the fusion break point were extracted and realigned to fusion contigs to improve interpretability of figures. The bottom panel shows the alignment of fusion supporting long reads.

Another fusion transcript, ENSG00000288891-MLT1A(LTR) had multiple reads cross the identified breakpoint in the heavy polysome fraction (**Supplementary Figure 5**). Interestingly, both the long read sequencing reads that initially identified the fusion transcript and the polysome sequencing reads identified the same SNP within the TE portion of the fusion transcript. This lends more support to the structure of the fusion transcript and that these are unique alignments from the same genomic origin. Another notable observation was the ZNF791-LTR2B(LTR) fusion transcript. This fusion transcript is expressed as two different isoforms, one originating from the canonical first exon of ZNF791 and one initiating from the second exon (**Figure 7**). Although both fusion isoforms associate with ribosomes, the isoform preserving the first exon was identified in the light polysome fraction suggesting association with multiple ribosomes while the isoform lacking the first exon was identified only in the monosome fraction. These data suggest that the transcriptional architecture at the 5′ end of gene–TE fusion isoforms can influence how far those isoforms progress into polysome loading. Future studies will be needed to identify whether the translation of these fusion transcripts generates a functional protein product and the functional impact they have.

## Discussion

Transposable elements (TEs) have long been recognized as powerful regulators of genome architecture and gene expression, influencing cellular identity across diverse biological contexts. In cancer, numerous studies have shown that TEs contribute to transcriptional rewiring by providing novel transcription factor binding sites, alternative promoters, and cryptic splice sites. However, the extent to which TEs shape the transcriptomic and translational identity of glioblastoma stem cells (GSCs), the self-renewing, tumor-propagating population within glioblastoma, has remained largely unexplored.

In this study, we integrated long-read sequencing with polysome profiling to systematically define how TEs influence RNA processing and translation in GSCs. We found that TE insertions frequently contribute to alternative splicing, leading to isoform switching within key oncogenic pathways such as p53 signaling and G2–M checkpoint regulation. Notably, isoforms containing TE sequences exhibited a significantly higher probability of altered ribosomal engagement (odds ratio: 2.54), suggesting that TE-associated splice events can modulate ribosomal association and potentially translational efficiency. Shorter TE-containing isoforms tended to accumulate in monosome fractions (odds ratio: 0.71), whereas longer isoforms were enriched in heavy polysomes (odds ratio: 1.26), indicating that TEs may influence the translational fate of transcripts through structural or regulatory effects on ribosome loading. Although isoform-specific proteomic validation will be required to determine whether these changes yield distinct protein isoforms, our findings reveal that TE-driven splicing represents a previously unappreciated layer of translational control in GSCs. Beyond their role in splicing regulation, we also sought to characterize TE exonization and gene-TE fusion events, which remain challenging to study with conventional short-read approaches due to the repetitive nature of TE sequences. To overcome this limitation, we developed FuTER (Fusion TE Reporter), a novel computational pipeline to identify TE incorporated fusion transcripts including those derived from novel TE insertions. FuTER leverages the full-length junctional information captured by long-read sequencing to detect gene–TE chimeric transcripts with high sensitivity and specificity, as confirmed using simulated libraries containing TE fusion transcripts. We further demonstrated that FuTER can identify previously unannotated TE insertions driving exonization events by applying it to publicly available reference genomes.

Applying FuTER to our GSC datasets, we identified 78 gene–TE exonization and long-range fusion events, including several involving cancer-relevant genes such as OS9. Manual inspection of transcript breakpoints and analysis of polysome fractions confirmed the authenticity and translational potential of several of these fusions. While our current validation set is limited by the lack of a comprehensive, experimentally verified reference library of TE fusion structures, our approach establishes a framework for future benchmarking. Developing such a curated resource will be critical for assessing and comparing emerging computational tools aimed at decoding TE-derived transcript diversity.

Although the functional impact of the identified TE fusion transcripts remains to be explored, their association with actively translating ribosomal fractions suggests potential protein-coding capacity. Follow-up proteomic and functional studies will be essential to elucidate whether these fusion transcripts produce novel protein isoforms or contribute to the malignant phenotype of GSCs.

Despite these limitations, our study provides the most comprehensive analysis to date of TE-mediated transcriptomic remodeling in glioblastoma stem cells. By integrating high-resolution long-read sequencing with polysome profiling, we reveal how TEs shape both the splicing and translational landscapes of these cells. Furthermore, FuTER represents a first-in-class, modular computational framework for the discovery and characterization of gene–TE fusion transcripts.

Together, these findings expand our understanding of TE-driven transcriptome complexity in cancer and provide a foundation for investigating how these elements contribute to tumor evolution, heterogeneity, and therapeutic resistance.

## Methods

### Cell culture

Primary GSC spheres were cultured form human glioma samples as previously described (55,56). In short, after obtaining informed written consent from patients in a completely deidentified manner at Geisinger Medical Center. Studies were performed in accordance with recognized ethical guidelines (Belmont Report). Cells were used after histological confirmation of glioblastoma diagnosis. Although in culture cells recapitulate the heterogeneity of tumors and contain multiple subtypes, GB2 and GB13 have been characterized as primarily proneural while GB24 is primarily mesenchymal and GB11 is the classical subtype (55,56).

Cells were used within 20 passages of being thawed out from liquid nitrogen storage and were cultured in a medium of 1X Neurobasal Medium (Fisher Scientific, 12-349-015), B27 serum-free supplement, minus Vitamin A (Fisher Scientific, 12587010), 100X Glutamax (Fisher Scientific, 35-050-061), 1 mg/mL Heparin (STEMCELL Technologies, 07980), 20 ng/mL epidermal growth factor (Peprotech, 100-47), 20 ng/mL bFGF (Peprotech, 100-18B). All GSC cultures are routinely tested for Mycoplasma contamination using the MycoSensor qPCR assay (Agilent, 302107). Cells were grown adhered to fibronectin (10 mg/mL, Millipore Sigma, FC0105MG) -coated plates (Thermo Scientific, 12-600-002).

Neural stem cells were obtained from NIH and grown adhered to cell culture dishes using Geltrex (Fisher Scientific, A1569601). They were grown in the same media as GSCs, supplemented with 1X NEAA MEM (Fisher Scientific, 11-140-050).

### RNA Collection and Sequencing

Total RNA was extracted using TRIzol™ Reagent (Thermo Fisher Scientific, 15-596-018) following the manufacturer’s protocol. GSCs were cultured as neurospheres until reaching approximately 80% confluency. After collection in TRIzol, cells were homogenized by thorough pipetting, and RNA was purified. Total RNA was evaluated for purity using a Nanodrop 2000 (ThermoFisher) and for integrity using an Agilent TapeStation 4150. To ensure high quality long read sequencing, samples were only utilized if the A260/A280 and A260/230 was > 1.9 and the RIN was > 8.5. RNA samples were prepared using Oxford Nanopore Technology’s cDNA-PCR Sequencing Kit V14 (SQK-PCS114) according to the manufacturer’s protocol. 50 ng of the finished library was loaded onto a R10.4.1 PromethION Flow Cell (FLO-PRO114M) and sequenced on a Promethion P2-Solo. Libraries were sequenced until at lead 30 million reads were collected to ensure deep coverage and help identify any low frequency fusion events.

### Polysome Sequencing

To determine whether any of the identified TE-Gene fusion transcripts are being actively translated, polysome sequencing of three GSC lines was performed. Cells were grown adhered to cell culture plates until they reached ∼70% confluence. At this point, cycloheximide (100 ug/mL, Fisher Scientific AAJ6690103) was added to the medium and cells were incubated for 15 min at 37degC.

Cells were collected using Accutase (Fisher Scientific, A1110501) and complete media and spun down in a centrifuge at 1000 rpm at 4 deg C for 5 min. The supernatant was removed, and cells were washed three times using pre chilled PBS containing 100 ug/mL cycloheximide. After the final wash, the supernatant was removed and the snap frozen samples were sent to Creative Biolabs for processing and sequencing. Raw FASTQ files were trimmed using trimmomatic (57). Transcript expression values were calculated using Salmon (58).

### Data Processing and Analysis

Basecalling of long-read sequencing data was performed using dorado (0.7.2+9ac85c6, *-emit-fastq -no-trim -min-qscore 7*). Output fastq files were filtered and trimmed of primer content using pychopper (2.7.10). Sequencing results were then aligned to the human genome (Gencode v44) using minimap2. We subsequently utilized Bambu (version 3.6.0), followed by SQANTI QC, ml-filter and ml-rescue modules (version 5.5.1) to obtain as high quality of a transcriptome as possible (38,39). We evaluated the resulting transcriptome using BUSCO (version 5.8.0), which identified all 11833 BUSCOs (only 1 of which was fragmented) (59). Differential expression analysis was performed using DESeq on gene counts (version 1.44.0) and lfcshrink apeglm was used to reduce noise in the fold change estimates (60,61). The processing pipeline described in more detail below was then used to identify gene-TE chimeric transcripts.

### Alternative Splicing Analysis

The bulk of the alternative splicing analysis was done in R (version 4.4.2) using a combination of different packages. Gviz (v1.48.0), ggplot2 (v3.5.2), and GenomicRanges (v1.56.2) were used throughout to generate figures and process data.

To identify alternative splicing patterns in our samples, we utilized IsoformSwitchAnalyzeR (ISA) (v 2.1.4). This approach allows us to investigate both alternative splicing as well as differential isoform usage (ie whether alternatively spliced isoforms of a gene are expressed at different levels between conditions). Internally, ISA utilizes DEXSeq (v1.50.0) to identify AS events. To identify TE involvement with alternative splicing (AS) events, we defined areas of interest (AOI) based on the borders of exons involved in AS. We extracted the location of the AS event and defined the AOI as the 150 bps into the intronic area. The width of the AOIs were defined based on previous studies that demonstrated TE insertions within 150 bps of a splice site are most likely to disrupt splicing processes (40,41). Next, we identified overlaps between these AOI and TEs in the genome to generate a list of what we refer to as TE+ isoforms.

To identify underlying polysome distribution patterns of TE+ isoforms, we extracted the isoform fraction of all TE+ isoforms and clustered the isoforms using kmeans ConsensusClusterPlus (v1.68.0) (*maxK = 10, pItem = 0.75, reps = 250, clusterAlgorithm = “km”).* The average isoform fraction was calculated within each cluster in each polysome fraction. A similar approach was used to identify underlying patterns in the fraction of a gene’s overall expression in the different polysome fractions. Rather than grouping all TE+ isoforms together in this analysis however, we calculated the fraction of individual gene’s expression that arises from TE+ isoforms versus TE- isoforms.

Finally, to calculate the relative polysome association of a gene’s isoforms, the polysomal isoform fraction was calculated. This was done by first calculating a gene’s expression across all polysome fractions (monosome, light and heavy polysome) by summing the TPM of all isoforms for a given gene across the three fractions. The TPM for the isoforms within each fraction was then divided by the gene’s total expression in order to calculate the polysomal isoform fraction.

### TE-Fusion Transcript Identification

Prior to sample analysis, a library of reference TE sequences was developed. First, TE sequences were identified using Repeat Masker (www.repeatmasker.org/, version 4.2.1*, -e rmblast -norna -s - species human*). Following the repeat masker documentation, simple repeats were manually removed from the output file and fragments were assembled using the one code to find them all script (62). A custom script was developed to merge the different output files into a gtf file. In order to avoid the most ancient TE sites, TE were filtered to retain elements that were less than 25% divergent from the consensus sequence and did not overlap with gene exons. The sequences of these sites were extracted, and potential cryptic splice sites were identified using SpliceAI (1.3.1) (46). The reference TE sequences were then appended to the human genome to create a combined reference genome.

To identify potential fusion transcripts, a method developed in CTAT-LR-Fusion was adapted. To more efficiently screen large libraries, reads were first aligned to the reference TE library and those that mapped to TE fragment were labeled as potential fusion transcripts. Reads without any TE sequences were not carried forward. TE containing reads were then realigned to the combined reference genome using ctat-minimap2, which generates full read alignments only for reads that map to multiple genomic locations (63). This generated a preliminary list of fusion candidates that were filtered for their percent identity to the genomic loci they map to. For the gene portion of potential fusion, a default minimum percent identity of 70% was required while for TEs this was reduced to 60% to account for increased divergence in their sequences. These preliminary fusions were then further filtered according to two main criteria. First, expression had to be above a default value of 0.75 FFPM (at least 0.75 fusion long reads per 1 million). Secondly, the fusion construct had to fulfill one of three breakpoint criteria:

1. Both the gene side of the fusion construct had to be within 50 base pairs of an exon boundary and the breakpoint within the TE had within 50 bp of a predicted splice site
2. One side is within 50 bp of an exon boundary while the other is within 1000 bp of a boundary and there are multiple reads supporting the fusion construct
3. The gene side is within 50 bp of an exon boundary while the TE is within 10000 bp of a predicted splice site and the fusion construct has at least an FFPM of 2 supporting the construct (in a library of 30 million reads, this requires 60 reads).

The pipeline then proceeded similarly to CTAT-LR-Fusion, annotating and filtering fusion candidates to exclude fusions that involved known red herrings. In the next phase of the pipeline, the gene and TE involved in the fusion were formed into a fusion contig. Reads that aligned to both members of the fusion construct were realigned to these contigs to accurately identify breakpoints. Extensive benchmarking of different minimap2 settings did not significantly change performance and underscored the importance of obtaining a high-quality reference TE library.

### Read Simulation

To test the performance of the pipeline, a library of simulated reads was generated. A modified version of Fusion Gene Simulator was used to create gene-TE chimeric transcripts (https://github.com/FusionSimulatorToolkit/FusionSimulatorToolkit/wiki). For fusion transcripts consisting of a gene fused to a TE, randomly selected gene’s exonic sequences were extracted from the first exon to an internal splice site. A random cryptic acceptor splice site was then selected and the TE’s sequence after this splice site was extracted and appended to the end of the gene. For TE-gene fusion transcripts, the same was performed in reverse. After generating 500 simulated gene-TE and TE-gene chimeras, these were duplicated five times to ensure deeper coverage. This set of fusions were subsequently combined with the 2,500 simulated gene-gene fusion transcripts developed by Haas and colleagues and all annotated transcripts (43). Reads were simulated using PBSIM3’s full length template simulation (*--strategy templ --method errhmm –errhmm data/ERRHMM-ONT-HQ.model --accuracy-mean 0.98*) (64). Reported benchmarking statistics were computed across three different sets of simulated reads.

### HCC1395 and HG002 Validation

The complete genome of HG002 was obtained from the genome in a bottle (GIAB) ftp website, along with the direct RNA sequencing files (65). The transposable element annotation was obtained from the UCSD Genome Browser (https://genome.ucsc.edu/cgi-bin/hgTrackUi?hgsid=3247758841_GCnJHAcauS9R7ay98EI5lzkZg5ZU&db=hub_4837794_HG002v1.1.MAT&c=chr3_MATERNAL&g=hub_4837794_HG002v1_1_MAT_repeatannots) (66). The HCC1395-BL sequencing files were obtained from the SRA using sratools (PRJNA934933) (67). The genome assembly for HCC1395 was obtained from the NCBI ftp site (https://ftp-trace.ncbi.nlm.nih.gov/ReferenceSamples/seqc/Somatic_Mutation_WG/assembly/) (68). Repeat masker was run on the HCC1395 assembly using the same settings as above. Reads from all samples were processed using the FuTER pipeline.

## DATA ACCESS

FuTER wil be publicly available on a GitHub repository (https://github.com/mpizzagalli777/FuTER), along with the TE reference library and all of the scripts used to generate figures for the paper. All sequencing data is available on GEO (GSE326354, GSE326349).

## Bibliography

1. Wells JN, Feschotte C. A Field Guide to Eukaryotic Transposable Elements. Annu Rev Genet. 2020 Nov 23;(54):539–61. doi:10.1146/annurev-genet-040620

2. Gebrie A. Transposable elements as essential elements in the control of gene expression. Mob DNA. 2023 Dec 1;14(1):9. doi:10.1186/S13100-023-00297-3 PubMed PMID: 37596675.

3. Lawson HA, Liang Y, Wang T. Transposable elements in mammalian chromatin organization. Nature Reviews Genetics 2023 24:10. 2023 Jun 7;24(10):712–23. doi:10.1038/s41576-023-00609-6 PubMed PMID: 37286742.

4. Klein SJ, O’neill RJ, Klein SJ, O’neill RJ. Transposable elements: genome innovation, chromosome diversity, and centromere conflict. Chromosome Research 2018 26:1. 2018 Jan 13;26(1):5–23. doi:10.1007/S10577-017-9569-5 PubMed PMID: 29332159.

5. Deniz Ö, Frost JM, Branco MR. Regulation of transposable elements by DNA modifications. Nature Reviews Genetics 2019 20:7. 2019 Mar 12;20(7):417–31. doi:10.1038/s41576-019-0106-6 PubMed PMID: 30867571.

6. Burns KH. Transposable elements in cancer. Nature Reviews Cancer 2017 17:7. 2017 Jun 9;17(7):415–24. doi:10.1038/nrc.2017.35 PubMed PMID: 28642606.

7. Liang Y, Qu X, Shah NM, Wang T. Towards targeting transposable elements for cancer therapy. Nature Reviews Cancer 2024 24:2. 2024 Jan 16;24(2):123–40. doi:10.1038/s41568-023-00653-8 PubMed PMID: 38228901.

8. Hedges DJ, Deininger PL. Inviting Instability: Transposable elements, Double-strand breaks, and the Maintenance of Genome Integrity. Mutat Res. 2006 Mar 1;616(1–2):46. doi:10.1016/J.MRFMMM.2006.11.021 PubMed PMID: 17157332.

9. Fueyo R, Judd J, Feschotte C, Wysocka J. Roles of transposable elements in the regulation of mammalian transcription. Nature Reviews Molecular Cell Biology 2022 23:7. 2022 Feb 28;23(7):481–97. doi:10.1038/s41580-022-00457-y PubMed PMID: 35228718.

10. Zhang Q, Pan J, Cong Y, Mao J. Transcriptional Regulation of Endogenous Retroviruses and Their Misregulation in Human Diseases. Int J Mol Sci. 2022 Sep 1;23(17). doi:10.3390/IJMS231710112 PubMed PMID: 36077510.

11. Drongitis D, Aniello F, Fucci L, Donizetti A. Roles of Transposable Elements in the Diierent Layers of Gene Expression Regulation. Int J Mol Sci. 2019 Nov 2;20(22):5755. doi:10.3390/IJMS20225755 PubMed PMID: 31731828.

12. Fuentes DR, Swigut T, Wysocka J. Systematic perturbation of retroviral LTRs reveals widespread long-range eiects on human gene regulation. Elife. 2018 Aug 2;7:e35989. doi:10.7554/ELIFE.35989 PubMed PMID: 30070637.

13. Schai LR, Mellinghoi IK. Glioblastoma and other Primary Brain Malignancies in Adults. A Review. JAMA. 2023 Feb 21;329(7):574. doi:10.1001/JAMA.2023.0023 PubMed PMID: 36809318.

14. Wen PY, Weller M, Lee EQ, Alexander BM, Barnholtz-Sloan JS, Barthel FP, et al. Glioblastoma in adults: a Society for Neuro-Oncology (SNO) and European Society of Neuro-Oncology (EANO) consensus review on current management and future directions. Neuro Oncol. 2020 Aug 1;22(8):1073. doi:10.1093/NEUONC/NOAA106 PubMed PMID: 32328653.

15. Chen J, Li Y, Yu TS, McKay RM, Burns DK, Kernie SG, et al. A restricted cell population propagates glioblastoma growth after chemotherapy. Nature 2012 488:7412. 2012 Aug 1;488(7412):522–6. doi:10.1038/nature11287 PubMed PMID: 22854781.

16. Bao S, Wu Q, McLendon RE, Hao Y, Shi Q, Hjelmeland AB, et al. Glioma stem cells promote radioresistance by preferential activation of the DNA damage response. Nature 2006 444:7120. 2006 Oct 18;444(7120):756–60. doi:10.1038/nature05236 PubMed PMID: 17051156.

17. Yuan Z, Yang Y, Zhang N, Soto C, Jiang X, An Z, et al. Human endogenous retroviruses in glioblastoma multiforme. Microorganisms. 2021 Apr 1;9(4). doi:10.3390/MICROORGANISMS9040764/S1

18. Bonté PE, Arribas YA, Merlotti A, Carrascal M, Zhang JV, Zueva E, et al. Single-cell RNA-seq-based proteogenomics identifies glioblastoma-specific transposable elements encoding HLA-I-presented peptides. Cell Rep. 2022 Jun 7;39(10). doi:10.1016/J.CELREP.2022.110916 PubMed PMID: 35675780.

19. Drews RM, Hernando B, Tarabichi M, Haase K, Lesluyes T, Smith PS, et al. A pan-cancer compendium of chromosomal instability. Nature 2022 606:7916. 2022 Jun 15;606(7916):976–83. doi:10.1038/s41586-022-04789-9 PubMed PMID: 35705807.

20. Mazzoleni A, Awuah WA, Sanker V, Bharadwaj HR, Aderinto N, Tan JK, et al. Chromosomal instability: a key driver in glioma pathogenesis and progression. European Journal of Medical Research 2024 29:1. 2024 Sep 4;29(1):1–17. doi:10.1186/S40001-024-02043-8 PubMed PMID: 39227895.

21. Hosea R, Hillary S, Naqvi S, Wu S, Kasim V. The two sides of chromosomal instability: drivers and brakes in cancer. Signal Transduction and Targeted Therapy 2024 9:1. 2024 Mar 29;9(1):1–30. doi:10.1038/s41392-024-01767-7 PubMed PMID: 38553459.

22. Dorney R, Dhungel BP, Rasko JEJ, Hebbard L, Schmitz U. Recent advances in cancer fusion transcript detection. Brief Bioinform. 2023 Jan 19;24(1):1–12. doi:10.1093/BIB/BBAC519 PubMed PMID: 36527429.

23. Gao Q, Liang WW, Foltz SM, Mutharasu G, Jayasinghe RG, Cao S, et al. Driver Fusions and Their Implications in the Development and Treatment of Human Cancers. Cell Rep. 2018 Apr 3;23(1):227–238.e3. doi:10.1016/J.CELREP.2018.03.050 PubMed PMID: 29617662.

24. Graham RP, Nair AA, Davila JI, Jin L, Jen J, Sukov WR, et al. Gastroblastoma harbors a recurrent somatic MALAT1-GLI1 fusion gene. Mod Pathol. 2017 Oct 1;30(10):1443–52. doi:10.1038/MODPATHOL.2017.68 PubMed PMID: 28731043.

25. Arribas YA, Baudon B, Rotival M, Suárez G, Bonté PE, Casas V, et al. Transposable element exonization generates a reservoir of evolving and functional protein isoforms. Cell. 2024 Dec 26;187(26):7603–7620.e22. doi:10.1016/J.CELL.2024.11.011 PubMed PMID: 39667937.

26. Burbage M, Rocañín-Arjó A, Baudon B, Arribas YA, Merlotti A, Rookhuizen DC, et al. Epigenetically controlled tumor antigens derived from splice junctions between exons and transposable elements. Sci Immunol. 2023 Feb 1;8(80). doi:10.1126/SCIIMMUNOL.ABM6360/SUPPL_FILE/SCIIMMUNOL.ABM6360_MDAR_REPRODUCIBILITY_CHECKLIST.PDF PubMed PMID: 36735776.

27. Merlotti A, Sadacca B, Arribas YA, Ngoma M, Burbage M, Goudot C, et al. Noncanonical splicing junctions between exons and transposable elements represent a source of immunogenic recurrent neo-antigens in patients with lung cancer. Sci Immunol. 2023 Feb 1;8(80). doi:10.1126/SCIIMMUNOL.ABM6359/SUPPL_FILE/SCIIMMUNOL.ABM6359_MDAR_REPRODUCIBILITY_CHECKLIST.PDF PubMed PMID: 36735774.

28. Mohammad T, Zolotovskaia MA, Suntsova M V., Buzdin AA. Cancer fusion transcripts with human non-coding RNAs. Front Oncol. 2024 Jun 11;14:1415801. doi:10.3389/FONC.2024.1415801/XML

29. Marasco LE, Kornblihtt AR. The physiology of alternative splicing. Nature Reviews Molecular Cell Biology 2022 24:4. 2022 Oct 13;24(4):242–54. doi:10.1038/s41580-022-00545-z PubMed PMID: 36229538.

30. Sibley CR, Blazquez L, Ule J. Lessons from non-canonical splicing. Nature Reviews Genetics 2016 17:7. 2016 May 31;17(7):407–21. doi:10.1038/nrg.2016.46 PubMed PMID: 27240813.

31. Saldi T, Riemondy K, Erickson B, Bentley DL. Alternative RNA structures formed during transcription depend on elongation rate and modify RNA processing. Mol Cell. 2021 Apr 15;81(8):1789–1801.e5. doi:10.1016/J.MOLCEL.2021.01.040 PubMed PMID: 33631106.

32. Shao Y, Chong W, Liu X, Xu Y, Zhang H, Xu Q, et al. Alternative splicing-derived intersectin1-L and intersectin1-S exert opposite function in glioma progression. Cell Death Dis. 2019 Jun 1;10(6). doi:10.1038/S41419-019-1668-0 PubMed PMID: 31160551.

33. Lan C, Zhang H, Wang K, Liu X, Zhao Y, Guo Z, et al. The alternative splicing of intersectin 1 regulated by PTBP1 promotes human glioma progression. Cell Death & Disease 2022 13:9. 2022 Sep 28;13(9):1–15. doi:10.1038/s41419-022-05238-1 PubMed PMID: 36171198.

34. Oliveira DS, Fablet M, Larue A, Vallier A, Carareto CMA, Rebollo R, et al. ChimeraTE: a pipeline to detect chimeric transcripts derived from genes and transposable elements. Nucleic Acids Res. 2023 Oct 13;51(18):9764–84. doi:10.1093/NAR/GKAD671

35. Shah NM, Jang HJ, Liang Y, Maeng JH, Tzeng SC, Wu A, et al. Pan-cancer analysis identifies tumor-specific antigens derived from transposable elements. Nature Genetics 2023 55:4. 2023 Mar 27;55(4):631–9. doi:10.1038/s41588-023-01349-3

36. Bolger I, Shaw R, Tam OH, Roque CG, Jackson CA, O’Neill K, et al. TDP-43 dysfunction leads to the accumulation of cryptic transposable element-derived exons, crypTEs, in iPSC derived neurons and ALS/FTD patient tissues. bioRxiv. 2026 Jan 9;2026.01.09.698641. doi:10.64898/2026.01.09.698641

37. Qin Q, Popic V, Wienand K, Yu H, White E, Khorgade A, et al. Accurate fusion transcript identification from long- and short-read isoform sequencing at bulk or single-cell resolution. Genome Res. 2025 Mar 14;35(4):967–86. doi:10.1101/GR.279200.124 PubMed PMID: 40086881.

38. Chen Y, Sim A, Wan YK, Yeo K, Lee JJX, Ling MH, et al. Context-aware transcript quantification from long-read RNA-seq data with Bambu. Nature Methods 2023 20:8. 2023 Jun 12;20(8):1187–95. doi:10.1038/s41592-023-01908-w PubMed PMID: 37308696.

39. Pardo-Palacios FJ, Arzalluz-Luque A, Kondratova L, Salguero P, Mestre-Tomás J, Amorín R, et al. SQANTI3: curation of long-read transcriptomes for accurate identification of known and novel isoforms. Nature Methods 2024 21:5. 2024 Mar 20;21(5):793–7. doi:10.1038/s41592-024-02229-2 PubMed PMID: 38509328.

40. Lev-Maor G, Ram O, Kim E, Sela N, Goren A, Levanon EY, et al. Intronic Alus Influence Alternative Splicing. PLoS Genet. 2008 Sep;4(9):e1000204. doi:10.1371/JOURNAL.PGEN.1000204 PubMed PMID: 18818740.

41. Zhang Y, Romanish MT, Mager DL. Distributions of Transposable Elements Reveal Hazardous Zones in Mammalian Introns. PLoS Comput Biol. 2011;7(5):e1002046. doi:10.1371/JOURNAL.PCBI.1002046 PubMed PMID: 21573203.

42. Raman P, Rominger MC, Young JM, Molaro A, Tsukiyama T, Malik HS. Novel Classes and Evolutionary Turnover of Histone H2B Variants in the Mammalian Germline. Mol Biol Evol. 2022 Feb 1;39(2):msac019. doi:10.1093/MOLBEV/MSAC019 PubMed PMID: 35099534.

43. Haas BJ, Dobin A, Li B, Stransky N, Pochet N, Regev A. Accuracy assessment of fusion transcript detection via read-mapping and de novo fusion transcript assembly-based methods. Genome Biol. 2019 Oct 21;20(1):1–16. doi:10.1186/S13059-019-1842-9/FIGURES/4 PubMed PMID: 31639029.

44. Zarnack K, König J, Tajnik M, Martincorena I, Eustermann S, Stévant I, et al. Direct competition between hnRNP C and U2AF65 protects the transcriptome from the exonization of Alu elements. Cell. 2013 Jan 31;152(3):453–66. doi:10.1016/J.CELL.2012.12.023 PubMed PMID: 23374342.

45. Attig J, Agostini F, Gooding C, Chakrabarti AM, Singh A, Haberman N, et al. Heteromeric RNP Assembly at LINEs Controls Lineage-Specific RNA Processing. Cell. 2018 Aug 23;174(5):1067–1081.e17. doi:10.1016/J.CELL.2018.07.001 PubMed PMID: 30078707.

46. Hershey JWB, Sonenberg N, Mathews MB. Principles of translational control: an overview. Cold Spring Harb Perspect Biol. 2012 Dec 1;4(12):a011528–a011528. doi:10.1101/CSHPERSPECT.A011528 PubMed PMID: 23209153.

47. Liang S, Bellato HM, Lorent J, Lupinacci FCS, Oertlin C, van Hoef V, et al. Polysome-profiling in small tissue samples. Nucleic Acids Res. 2018 Jan 9;46(1):e3–e3. doi:10.1093/NAR/GKX940 PubMed PMID: 29069469.

48. Chassé H, Boulben S, Costache V, Cormier P, Morales J. Analysis of translation using polysome profiling. Nucleic Acids Res. 2016 Feb 17;45(3):e15. doi:10.1093/NAR/GKW907 PubMed PMID: 28180329.

49. Ruggero D. Translational control in cancer etiology. Cold Spring Harb Perspect Biol. 2013;5(2). doi:10.1101/CSHPERSPECT.A012336 PubMed PMID: 22767671.

50. Heyer EE, Correspondence MJM. Redefining the Translational Status of 80S Monosomes. Cell. 2016;164:757–69. doi:10.1016/j.cell.2016.01.003

51. Gandin V, Masvidal L, Hulea L, Gravel SP, Cargnello M, McLaughlan S, et al. NanoCAGE reveals 5’ UTR features that define specific modes of translation of functionally related MTOR-sensitive mRNAs. Genome Res. 2016 May 1;26(5):636–48. doi:10.1101/GR.197566.115/-/DC1 PubMed PMID: 26984228.

52. Haas BJ, Dobin A, Ghandi M, Van Arsdale A, Tickle T, Robinson JT, et al. Targeted in silico characterization of fusion transcripts in tumor and normal tissues via FusionInspector. Cell Reports Methods. 2023 May 22;3(5):100467. doi:10.1016/J.CRMETH.2023.100467 PubMed PMID: 37323575.

53. Zepecki JP, Snyder KM, Moreno MM, Fajardo E, Fiser A, Ness J, et al. Regulation of human glioma cell migration, tumor growth, and stemness gene expression using a Lck targeted inhibitor. Oncogene 2018 38:10. 2018 Oct 23;38(10):1734–50. doi:10.1038/s41388-018-0546-z PubMed PMID: 30353164.

54. Guetta-Terrier C, Karambizi D, Akosman B, Zepecki JP, Chen JS, Kamle S, et al. Chi3l1 Is a Modulator of Glioma Stem Cell States and a Therapeutic Target in Glioblastoma. Cancer Res. 2023 Jun 6;83(12):1984. doi:10.1158/0008-5472.CAN-21-3629 PubMed PMID: 37101376.

55. Bolger AM, Lohse M, Usadel B. Trimmomatic: a flexible trimmer for Illumina sequence data. Bioinformatics. 2014 Aug 1;30(15):2114. doi:10.1093/BIOINFORMATICS/BTU170 PubMed PMID: 24695404.

56. Patro R, Duggal G, Love MI, Irizarry RA, Kingsford C. Salmon provides fast and bias-aware quantification of transcript expression. Nature Methods 2017 14:4. 2017 Mar 6;14(4):417–9. doi:10.1038/nmeth.4197 PubMed PMID: 28263959.

57. Simão FA, Waterhouse RM, Ioannidis P, Kriventseva E V., Zdobnov EM. BUSCO: assessing genome assembly and annotation completeness with single-copy orthologs. Bioinformatics. 2015 Oct 1;31(19):3210–2. doi:10.1093/BIOINFORMATICS/BTV351 PubMed PMID: 26059717.

58. Love MI, Huber W, Anders S. Moderated estimation of fold change and dispersion for RNA-seq data with DESeq2. Genome Biol. 2014 Dec 5;15(12):1–21. doi:10.1186/S13059-014-0550-8/FIGURES/9 PubMed PMID: 25516281.

59. Zhu A, Ibrahim JG, Love MI. Heavy-tailed prior distributions for sequence count data: removing the noise and preserving large diierences. Bioinformatics. 2019 Jun 15;35(12):2084–92. doi:10.1093/BIOINFORMATICS/BTY895 PubMed PMID: 30395178.

60. Bailly-Bechet M, Haudry A, Lerat E. “One code to find them all”: A perl tool to conveniently parse RepeatMasker output files. Mob DNA. 2014 May 1;5(1):1–15. doi:10.1186/1759-8753-5-13/TABLES/3

61. Jaganathan K, Kyriazopoulou Panagiotopoulou S, McRae JF, Darbandi SF, Knowles D, Li YI, et al. Predicting Splicing from Primary Sequence with Deep Learning. Cell. 2019 Jan 24;176(3):535–548.e24. doi:10.1016/J.CELL.2018.12.015 PubMed PMID: 30661751.

62. Li H. Minimap2: pairwise alignment for nucleotide sequences. Bioinformatics. 2018 Sep 15;34(18):3094–100. doi:10.1093/BIOINFORMATICS/BTY191 PubMed PMID: 29750242.

63. Ono Y, Hamada M, Asai K. PBSIM3: a simulator for all types of PacBio and ONT long reads. NAR Genom Bioinform. 2022 Oct 6;4(4). doi:10.1093/NARGAB/LQAC092

64. Hansen NF, Dwarshuis N, Ji HJ, Rhie A, Loucks H, Logsdon GA, et al. A complete diploid human genome benchmark for personalized genomics. bioRxiv. 2025 Sep 21;2025.09.21.677443. doi:10.1101/2025.09.21.677443

65. Hoyt SJ, Storer JM, Hartley GA, Grady PGS, Gershman A, de Lima LG, et al. From telomere to telomere: The transcriptional and epigenetic state of human repeat elements. Science (1979). 2022 Apr 1;376(6588). doi:10.1126/SCIENCE.ABK3112 PubMed PMID: 35357925.

66. Cotto KC, Feng YY, Ramu A, Richters M, Freshour SL, Skidmore ZL, et al. Integrated analysis of genomic and transcriptomic data for the discovery of splice-associated variants in cancer. Nature Communications 2023 14:1. 2023 Mar 22;14(1):1–18. doi:10.1038/s41467-023-37266-6 PubMed PMID: 36949070.

67. Xiao C, Chen Z, Chen W, Padilla C, Colgan M, Wu W, et al. Personalized genome assembly for accurate cancer somatic mutation discovery using tumor-normal paired reference samples. Genome Biol. 2022 Dec 1;23(1):1–34. doi:10.1186/S13059-022-02803-X/TABLES/2 PubMed PMID: 36352452.

