## Supplemental Figure and Tables for "A long-read RNA sequencing and polysome profiling framework reveals transposable element–driven transcript diversity and translational rewiring in glioblastoma"

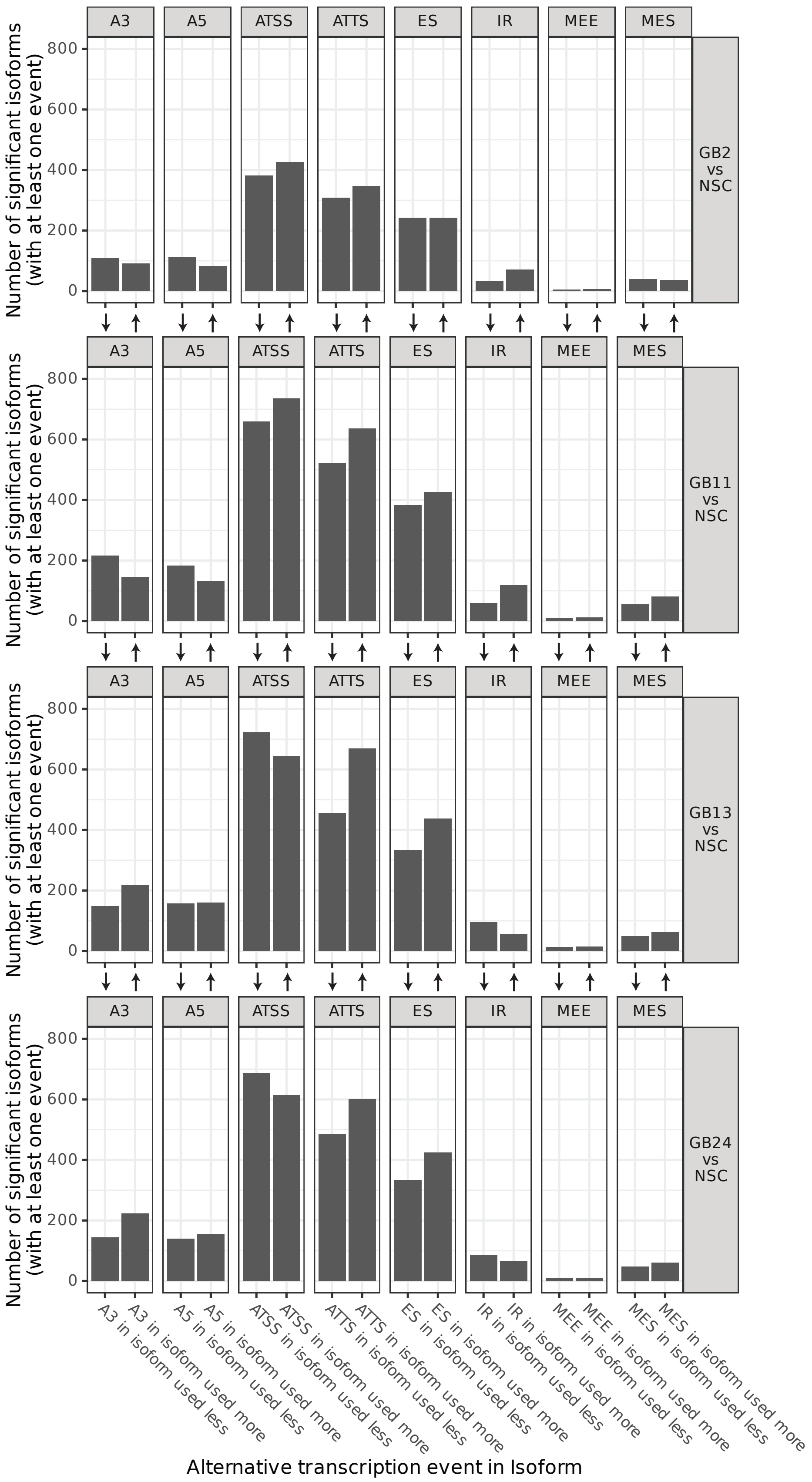
Number of significant isoforms (with at least one event)
A3
A5
ATSS
ATTS
ES
IR
MEE
MES
GB2 vs NSC
GB11 vs NSC
GB13 vs NSC
GB24 vs NSC
0
200
400
600
800
A3 in isoform used less
A3 in isoform used more
A5 in isoform used less
A5 in isoform used more
ATSS in isoform used less
ATSS in isoform used more
ATTS in isoform used less
ATTS in isoform used more
ES in isoform used less
ES in isoform used more
IR in isoform used less
IR in isoform used more
MEE in isoform used less
MEE in isoform used more
MES in isoform used less
MES in isoform used more
Alternative transcription event in Isoform

**Supplementary Figure 1: Summary of Global Splicing Changes Between Indicated GSC and NSCs.** Upward facing arrows indicate that the corresponding AS event (ie A3, A5, ES, etc) is used more in GB24 vs NCS. Downward arrow indicates the opposite. Full labels are located at the bottom of the series of plots. A3: Alternative 3' Splice Site; A5: Alternative 5' Splice Site; ATSS: Alternative Transcription Start Site; ATTS: Alternative Transcription Termination Site; ES: Exon Skipping; IR: Intron Retention; MEE: Mutually Exclusive Exons; MES: Mutual Exon Skipping

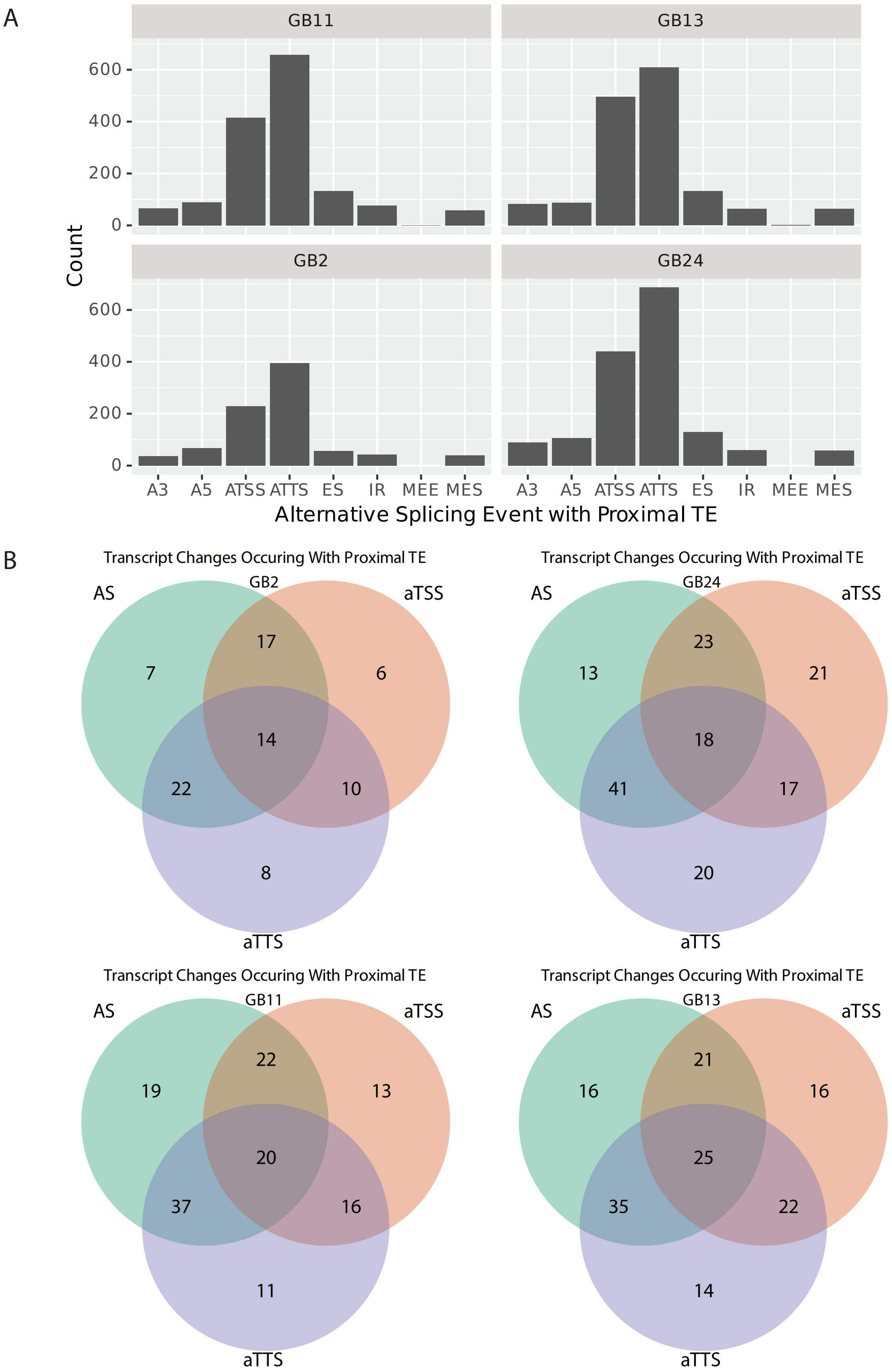
A
GB11
GB13
GB2
GB24
Count
600
400
200
0
A3
A5
ATSS
ATTS
ES
IR
MEE
MES
Alternative Splicing Event with Proximal TE
B
Transcript Changes Occuring With Proximal TE
GB2
AS
aTSS
aTTS
7
17
6
14
22
10
8
Transcript Changes Occuring With Proximal TE
GB24
AS
aTSS
aTTS
13
23
21
18
41
17
20
Transcript Changes Occuring With Proximal TE
GB11
AS
aTSS
aTTS
19
22
13
20
37
16
11
Transcript Changes Occuring With Proximal TE
GB13
AS
aTSS
aTTS
16
21
16
25
35
22
14

**Supplementary Figure 2: (A)** A breakdown of the types of transcriptomic changes that arise from an AS event with a proximal TE. **(B)** Venn diagrams showing the structural changes and splicing differences that arise from an AS event with a proximal TE that results in significant isoform switching.

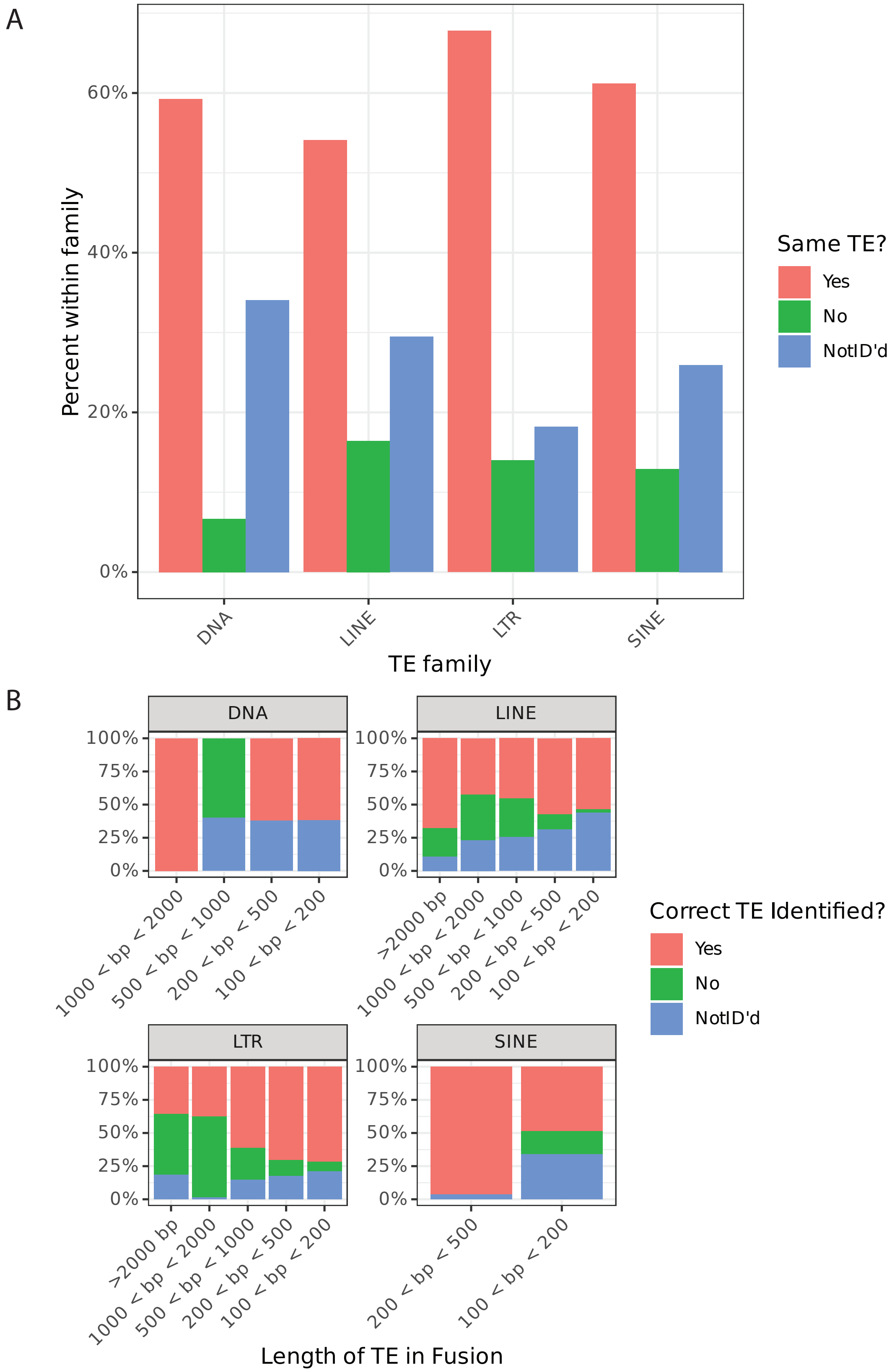
A
Percent within family
0%
20%
40%
60%
DNA
LINE
LTR
SINE
TE family
Same TE?
Yes
No
NotID'd
B
DNA
LINE
LTR
SINE
100%
75%
50%
25%
0%
>2000 bp
1000 < bp < 2000
500 < bp < 1000
200 < bp < 500
100 < bp < 200
Length of TE in Fusion
Correct TE Identified?
Yes
No
NotID'd

**Supplementary Figure 3: Performance of FuTER varies between families (A)** Performance of FuTER varies between families of TEs, with LTRs demonstrating the highest rate of identification (68%) while LINEs were correctly identified at the lowest rate (54%). Interestingly, Class I DNA TEs had the highest rate of non-identification and the lowest rate of incorrectly assigned TEs. This suggests that FuTER struggles to align DNA TEs. **(B)** The length of the TE partner in the fusion significantly impacts the performance of FuTER as well. Longer elements are aligned at a higher rate, although this does not always correlate with an increase in precision.

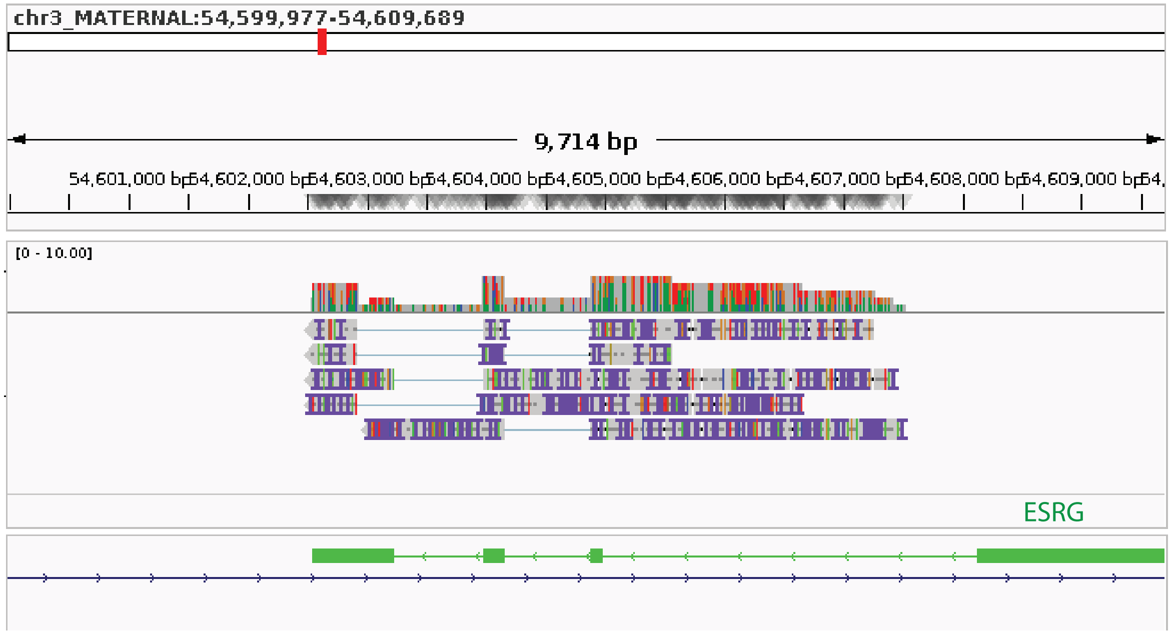


**Supplementary Figure 4:** FuTER identified a TE fusion involving ESRG in HG002. Manual inspection of the fusion demonstrated a significant number of insertions and deletions, indicating potential poor read quality or mapping giving rise to this false positive. This also demonstrated the importance of accurate sequencing data. These reads were generated using direct RNA sequencing, which has lagged behind cDNA based long read RNA sequencing in terms of base calling accuracy, although the approach does negate any risk of fold back or RT artifacts.

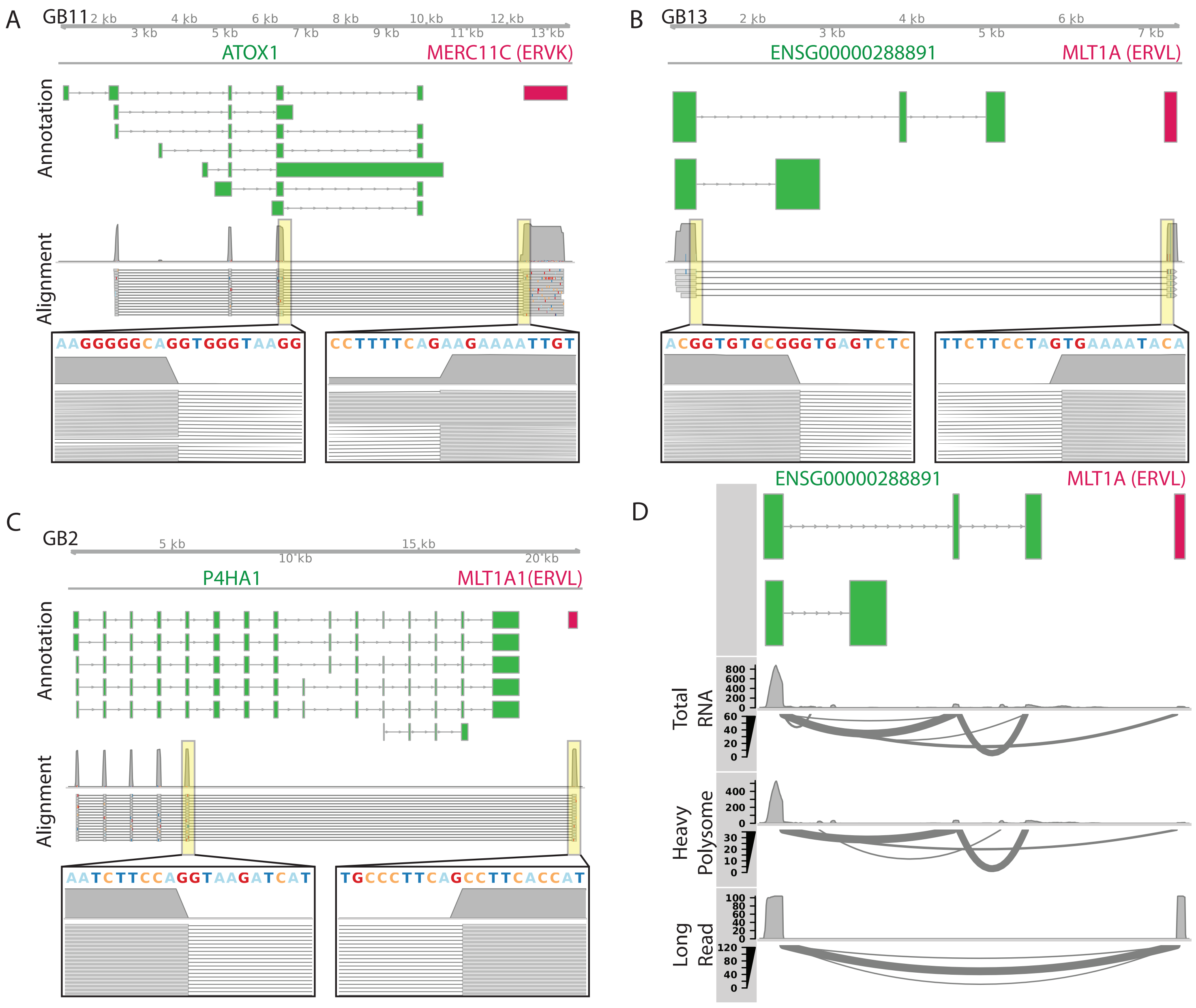

A
GB11
ATOX1
MERC11C (ERVK)
Annotation
Alignment
AAGGGGGCAGGTGGGTAAGG
CCTTTTCAGAAGAAAATTGT
B
GB13
ENSG00000288891
MLT1A (ERVL)
ACGGTGTGCGGGTGAGTCTC
TTCTTCCTAGTGAAAATACA
C
GB2
P4HA1
MLT1A1(ERVL)
Annotation
Alignment
AATCTTCCAGGTAAGATCAT
TGCCCTTCAGCCTTCACCAT
D
ENSG00000288891
MLT1A (ERVL)
Total RNA
Heavy Polysome
Long Read

**Supplementary Figure 5.** Further representative images showing numerous reads crossing the breakpoint of FuTER identified fusion transcripts. **A-C** show the fusion transcript structure in the top panel and reads that cleanly cross the breakpoint in the bottom panel (Alignment). Inserts show the exact sequences surrounding the breakpoint, which typically have consensus splice sites. **Figure D** incorporates the polysome profiling data for the fusion transcript arising from ENSG00000288891 and MLT1A (ERVL), which validated the fusion in an independently generated data set.

| Sample | Reads | Est Bases |
|---|---|---|
| GB2 1 | 37.82 M | 54.59 Gb |
| GB2 2 | 69.15 M | 90.09 Gb |
| GB2 3 | 38.83 M | 44.66 Gb |
| GB24 1 | 43.55 M | 47.39 Gb |
| GB24 2 | 56.75 M | 66.09 Gb |
| GB24 3 | 39.72 M | 42.01 Gb |
| GB11 1 | 40.62 M | 47.72 Gb |
| GB11 2 | 41.86 M | 54.79 Gb |
| GB11 3 | 32.83 M | 44.52 Gb |
| GB13 1 | 50.99 M | 55.88 Gb |
| GB13 2 | 36.38 M | 52.12 Gb |
| GB13 3 | 42.65 M | 52.67 Gb |
| H9 1 | 36.52 M | 45.01 Gb |
| H9 2 | 38.91 M | 50.31 Gb |
| H9 3 | 48.62 M | 53.11 GB |
| H14 1 | 38.81 M | 41.08 Gb |
| H14 2 | 38.13 M | 43.77 Gb |
| H14 3 | 33.71 M | 41.75 Gb |

**Supplementary Table 1:** Long Read Sequencing Library Information

| TE Family | TE Subfamily | Count |
|---|---|---|
| **DNA** | **TOTAL** | **224506** |
| DNA | Crypton | 121 |
| DNA | Crypton-A | 45 |
| DNA | Kolobok | 145 |
| DNA | MULE-MuDR | 1571 |
| DNA | Merlin | 121 |
| DNA | PIF-Harbinger | 114 |
| DNA | PiggyBac | 756 |
| DNA | RC | 381 |
| DNA | TcMar | 46 |
| DNA | TcMar-Marine | 6616 |
| DNA | TcMar-Pogo | 15 |
| DNA | TcMar-Tc1 | 370 |
| DNA | TcMar-Tc2 | 3345 |
| DNA | TcMar-Tigger | 53260 |
| DNA | hAT | 919 |
| DNA | hAT-Ac | 1070 |
| DNA | hAT-Blackjack | 8799 |
| DNA | hAT-Charlie | 121745 |
| DNA | hAT-Tag1 | 416 |
| DNA | hAT-Tip100 | 21806 |
| DNA | hAT-hAT19 | 13 |
| DNA | NA | 2832 |
| **LINE** | **TOTAL** | **345180** |
| LINE | CR1 | 6361 |
| LINE | Dong-R4 | 81 |
| LINE | I-Jockey | 35 |
| LINE | L1 | 285564 |
| LINE | L1-Tx1 | 14 |
| LINE | L2 | 50868 |
| LINE | RTE-BovB | 649 |
| LINE | RTE-X | 1608 |
| **LTR** | **TOTAL** | **298734** |
| LTR | DIRS | 2607 |
| LTR | ERV1 | 73645 |
| LTR | ERVK | 2370 |
| LTR | ERVL | 48888 |
| LTR | ERVL-MaLR | 164932 |
| LTR | Gypsy | 3452 |
| LTR | NA | 2840 |
| **SINE** | **TOTAL** | **732842** |
| SINE | 5S-Deu-L2 | 570 |
| SINE | Alu | 624862 |
| SINE | MIR | 106804 |
| SINE | tRNA-Deu | 112 |
| SINE | tRNA-Deu-L2 | 19 |
| SINE | tRNA-RTE | 475 |

**Supplementary Table 2:** Types of TEs included in Reference Set

| Type of Fusion | Count |
|---|---|
| GENE | 252835 |
| GENE-GENE | 12500 |
| GENE-TE | 1495 |
| TE-GENE | 1005 |

| TE Family | Count |
|---|---|
| DNA | 45 |
| LINE | 820 |
| LTR | 1440 |
| SINE | 195 |

**Supplementary Table 3:** Simulated Fusions represent the four major families of TEs
